# Polygenic adaptation, clonal interference, and the evolution of mutators in experimental *Pseudomonas aeruginosa* populations

**DOI:** 10.1101/2021.01.14.426720

**Authors:** Katrina B. Harris, Kenneth M. Flynn, Vaughn S. Cooper

## Abstract

In bacterial populations, switches in lifestyle from motile, planktonic growth to surface-grown biofilm is associated with persistence in both infections and non-clinical biofilms. Studies have identified the first steps of adaptation to biofilm growth but have yet to replicate the extensive genetic diversity observed in chronic infections or in the natural environment. We conducted a 90-day long evolution experiment with *Pseudomonas aeruginosa* PA14 in growth media that promotes biofilm formation in either planktonic culture or in a biofilm bead model. Surprisingly, all populations evolved extensive genetic diversity with hundreds of mutations maintained at intermediate frequencies, while fixation events were rare. Instead of the expected few beneficial mutations rising in frequency through successive sweeps, we observe a remarkable 40 genes with parallel mutations spanning both environments and often on coexisting genotypes within a population. Additionally, the evolution of mutator genotypes (*mutS* or *mutL* mutator alleles) that rise to high frequencies in as little as 25 days contribute to the extensive genetic variation and strong clonal interference. Parallelism in several transporters (including *pitA, pntB, nosD*, and *pchF*) indicate probable adaptation to the arginine media that becomes highly alkaline during growth. Further, genes involved in signal transduction (including *gacS, aer2, bdlA, and* PA14_71750) reflect likely adaptations to biofilm-inducing conditions. This experiment shows how extensive genetic and phenotypic diversity can arise and persist in microbial populations despite strong selection that would normally purge diversity.

**Importance:** How biodiversity arises and is maintained in clonally reproducing organisms like microbes remains unclear. Many models presume that beneficial genotypes will outgrow others and purge variation via selective sweeps. Environmental structure like biofilms may oppose this process and preserve variation. We tested this hypothesis by evolving *P. aeruginosa* populations in biofilm-promoting media for three months and found both adaptation and diversification that were mostly uninterrupted by fixation events that eliminate diversity. Genetic variation tended to be greater in lines grown using a bead model of biofilm growth but many lineages also persisted in planktonic lines. Convergent evolution affecting dozens of genes indicates that selection acted on a wide variety of traits to improve fitness, causing many adapting lineages to co-occur and persist. This result demonstrates that some environments may expose a large fraction of the genome to selection and select for many adaptations at once, causing enduring diversity.

## Introduction

Bacterial populations inhabit countless environments along a continuum of spatial structure, ranging from a well-mixed liquid to rugged, solid surfaces. Growth on surfaces is associated with biofilm production that in turn generates nutrient and oxygen gradients (Stoodley et al. 2002; Dietrich et al. 2013), but so too does the metabolic activities of neighboring cells. Life in confined spaces results in varied levels of nutrients, waste, and signaling molecules that may alter selective forces on parts of the population and create novel ecological opportunities (Poltak and Cooper 2011). These physical differences are hypothesized to select for mutations in biofilm populations distinct from well mixed cultures. Experimental microbial evolution (EME) studies have found that biofilm populations select for different mutations than those seen in planktonic growth, and the greater environmental structure of the biofilm produces increased genetic diversity (Rainey and Travisano 1998; Boles et al. 2004; Habets et al. 2006; Wong et al. 2012; Traverse et al. 2013; Flynn et al. 2016; Santos-Lopez et al. 2019). Despite this process of diversification, replicate populations propagated in both biofilm and planktonic conditions exhibit high levels of both phenotypic and genetic parallelism, suggesting some measure of predictability within the same environment (Wong et al. 2012; Tognon et al. 2017; Yen and Papin 2017; Sanz-García et al. 2018; Turner et al. 2018). We still have much to learn about how the biofilm life cycle influences evolutionary dynamics and processes, including the relative roles of mutation and selection, and whether biofilm growth becomes the dominant selective force relative to other stresses like nutrient limitation or external toxins.

*Pseudomonas aeruginosa* is an opportunistic pathogen found in soil and water and is known for its ability to thrive in numerous environments. *P. aeruginosa* is a highly studied pathogen due to its association with poor outcomes in clinical settings when it forms biofilm infections within patients (Eichner et al. 2014; Feliziani, Marvig, Luján, Moyano, Di Rienzo, et al. 2014; Schick and Kassen 2018; Gloag et al. 2019). Adaptation to biofilm growth has been indicated as a cause of infection persistence in numerous studies (Parsek and Singh 2003; Periasamy and Kolenbrander 2009; Yildiz and Visick 2009). This adaptation manifests as important colony phenotypes such as mucoidy and rugose small colony variants (RSCVs) as well as the loss of virulence factor production and altered cell surface virulence determinants (Ashish et al. 2013; Kim et al. 2014). Several studies have been performed to identify some of the first steps of adaptation to a biofilm environment but these studies have yet to replicate the high levels of genetic diversity often seen in chronic biofilm-associated infections, such as those of the airways of cystic fibrosis patients (Lieberman et al. 2011; Hogardt and Heesemann 2013; Marvig et al. 2015; Kruczek et al. 2016; Schick and Kassen 2018; Gloag et al. 2019). We previously conducted a long-term evolution experiment over 90 days of propagation (∼600 generations) to explore how the population-genetic dynamics of *P. aeruginosa* adaptation differ in a model of the biofilm life cycle compared to a well-mixed environment (Flynn et al. 2016). Major findings included that biofilm populations evolved greater phenotypic and ecological diversity than planktonic populations, and some of these phenotypes were linked to probable biofilm adaptations. The different colony types that evolved within populations represented distinct growth strategies that were most productive in mixture, suggesting a process of niche partitioning and facilitation among lineages (Flynn et al. 2016). This is strikingly opposed to the phenotypic parallelism observed in previous studies(Rainey and Travisano 1998; Traverse et al. 2013). A preliminary survey of the genomes of these lineages revealed that this diversity was in part caused by the evolution of *mutS* and *mutL* genotypes that increased mutation rate, a genotype also observed in many chronic infections of *P. aeruginosa* (Ciofu et al. 2010; Macià et al. 2011; Warren et al. 2011; López-Causapé et al. 2013). Here, we use longitudinal whole-population genome sequencing (WPGS) to study the underlying evolutionary dynamics of this experiment at high resolution. Some genes that experienced mutational parallelism contribute to arginine metabolism, the sole carbon and nitrogen source in these experiments. More surprising, in light of the duration of the experiment and the strength of selection in these populations, few mutations actually fixed in any of the six populations and much more genetic diversity was preserved than has been seen in previous evolution experiments. We discuss potential evolutionary and phenotypic causes for this maintenance of such levels of high genetic diversity.

## Results

### Experimental evolution

Six populations of *P. aeruginosa* strain PA14 (PA), a strain already proficient in forming biofilm, were propagated for 90 days (approximately 600 generations, or ∼6.6 generations per day) to examine the population-genetic dynamics of prolonged growth in a biofilm life cycle. Bacteria that attach to a polystyrene bead are transferred daily to a new tube containing a fresh bead (populations B1, B2, B3), and compared to passages of serial 1:100 dilutions (planktonic populations P1, P2, P3) as described previously (Poltak and Cooper 2011; Flynn et al. 2016) (Figure S1). Biofilm and planktonic populations were designed to ensure a similar transfer size and number of generations per day, as reported previously(Flynn et al. 2016). The culture medium was M63 media containing arginine as the sole carbon and nitrogen source and supplemented with 25 μM iron (see methods), a combination which has been shown to promote biofilm production in *Pseudomonas* species (Bernier et al. 2011). We hypothesized that this media would allow us to bypass many known first-step adaptive mutations, and instead allow us to look at the subsequent steps of adaptation that are less known. We sought to study these population wide adaptations at both phenotypic and genetic levels.

After 90 days, all six populations became vastly more fit than ancestral PA14, with selective coefficients 10-fold greater (*r* between 3 and 5; see methods for selection calculation) than the fitness gains observed in other evolution experiments of similar duration (Lenski et al. 1991; Rainey and Travisano 1998; Barrett et al. 2005; Ellis et al. 2012; Wong et al. 2012) (Figure S2, Table S1; see methods). Five populations increased their maximum growth rate (Vmax; Figure S3a) and five populations evolved to become less motile than the ancestor (Figure S3b). Despite selection to attach to a plastic bead each day, presumably by producing biofilm, only the B3 population evolved higher biofilm production than the ancestor as measured by the standard crystal violet assay. In contrast, four populations, including bead populations B1 and B2, evolved lower biofilm production (Figure S3c). This result may reflect the high starting biofilm production of PA14 in these conditions being difficult to improve. We performed a principal component analysis on these three phenotypes (Figure 1a) and found that evolved populations were distinguished from the ancestor but not clearly separated by treatment.

**Figure 1.**
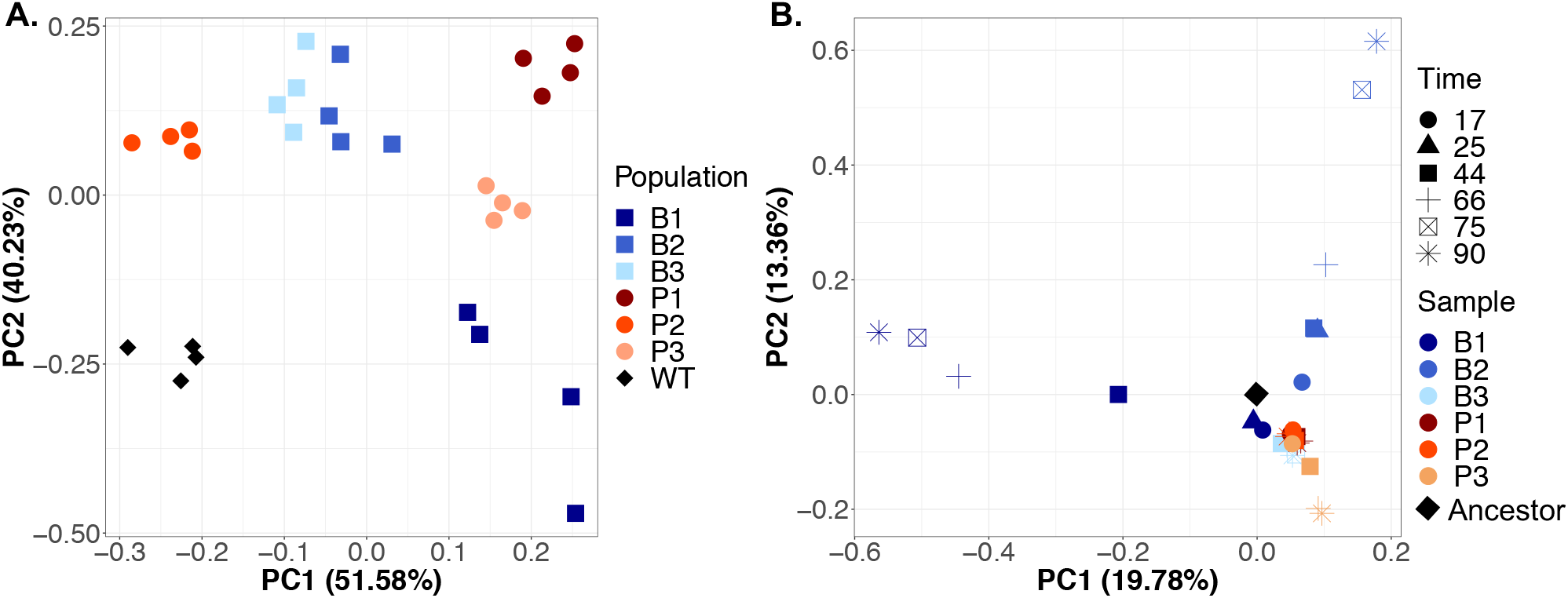
Divergent phenotypes linked to fitness and genotypes distinguish each evolved population after 90 days of passage. (A) PCA of evolved phenotypes including biofilm production, maximum growth rate (Vmax), and swimming motility (n = 4 for each population). Blue squares are biofilm populations, red circles are planktonic populations, and the ancestor is indicated by black diamond(s). The first three components explain 51.58%, 40.23%, and 8.19% of variance. (B) PCA of mutation identities and frequencies at Days 17, 25, 44, 66, 75, and 90. Replicate data points for each population are indicated in color. Colors are the same as in A, but shape indicates time through the experiment. The top four components explain 19.78%, 13.36%, 9.31%, and 6.5% of the variance.

### High genetic diversity arose and persisted in all populations

We used longitudinal WPGS to characterize the genetic diversity and evolutionary dynamics of each lineage with an average depth of 617.1± 142.3 reads per sample. A total of 145.7 ± 55 mutations were detected per population over the course of the 90-day time period (see methods for filtering criteria). Biofilm populations accumulated more mutations (535, with 382 found at day 90) over the 90 days than planktonic populations (339, with 201 found at day 90; Table 1). This number of mutations is remarkable in comparison to previous evolution experiments. For example, *Burkholderia cenocepacia* populations were propagated using the same bead model, for twice the transfer days, and 37 mutations were identified in the best-studied population (Traverse et al. 2013). Biofilm populations were sampled at two more time points than planktonic allowing for a more accurate representation of their complex dynamics, discussed more below.

**Table 1.** Summarized mutation statistics from all six populations following 90 days of experimental evolution.

|  |  |
| --- | --- |
| **Total cumulative mutations** | **874** |
| Biofilm only | 535 |
| Planktonic only | 339 |
| **Total mutations at day 90** | **583** |
| Biofilm only | 382 |
| Planktonic only | 201 |
| **Total fixed mutations** | **53** |
| Biofilm only | 48 |
| Planktonic only | 5 |
| **Total fixation events** | **10** |

A known phenomenon associated with positive selection is the increase in nonsynonymous (NS) mutations over synonymous (S) mutations (Kryazhimskiy and Plotkin 2008; Lieberman et al. 2011). In following with this trend, the normalized NS/S ratio for planktonic populations averaged 1.73 and for biofilm populations 2.34. A total of 31 indels were also identified in the six evolved populations (Table S2, Table S3) while, interestingly, no large structural variants were detected. A large number of intergenic mutations were also identified (233 mutations, or 27.5%), many in likely promoter or terminator regions. The importance of intergenic mutations still remains unclear, however recent work has implicated intergenic regulatory regions as critical in *Pseudomonas aeruginosa* evolution in the laboratory and during infections (Feliziani, Marvig, Luján, Moyano, Rienzo, et al. 2014; Khademi et al. 2019).

We hypothesized that the structured biofilm environment would maintain greater genetic diversity than mixed liquid culture, but this ecological process is potentially at odds with the diversity-purging effect of selection on beneficial mutations. For example, a mutation that generally improves growth would be expected to sweep and eliminate genetic variation, even though the biofilm growth is predicted to increase the probability of coexisting subpopulations (Habets et al. 2006; Traverse et al. 2013; Martin et al. 2016). Planktonic populations would also be subject to similar effects of selective sweeps but their admixture might limit the cooccurrence of contending mutations (Lang et al. 2013). To summarize, comparing populations at a given time point might not adequately capture the longitudinal ecological and evolutionary dynamics affecting diversity. For this reason, we sampled biofilm populations at six time points (17, 25, 44, 66, 75, and 90 days) and planktonic populations at four time points (17, 44, 66, and 90 days) and focused on temporal changes in all mutations supported by three reads on each of both strands.

The nucleotide diversity of biofilm populations tended to be greater at the end of the experiment than after only 17 transfers (day 17 vs day 90: p = 0.0478, t = 2.821, df = 4 via two tailed t-test). Planktonic populations continued to select for new mutations throughout the experiment, but they did not become more diverse by day 90 than at day 17 (p = 0.239, t = 1.384, df = 4 via two tailed t-test), indicating a process where new mutations were displacing older ones. Biofilm populations also tended toward greater diversity than planktonic populations later in the experiment, albeit not significantly (planktonic vs biofilm, day 90, p = 0.0749, t = 2.394, df = 4 via two tailed t-test; Figure S4a). These findings are consistent with the greater morphological variation in biofilm populations than planktonic ones that we reported previously (Flynn et al. 2016).

A major cause of the high genetic diversity is the prevalence of mutator alleles, mutations that cause a genome-wide increase in mutation rate, in four populations. In all biofilm populations and one planktonic population (P3) NS mutations in the DNA mismatch repair (MMR) genes *mutS* and *mutL* became frequent or fixed. Two populations, B1 and B2, independently evolved the same *mutS* mutation (T112P) that has been shown to lower affinity for heteroduplex DNA in *E. coli*(Junop et al. 2003) and results in a 116-fold increase in mutation rate in isogenic mutants (Figure S5). Interestingly, this T112P mutation occurs within a homopolymeric region that may be a mutational hotspot. Genotypes containing these mutations fix in populations B1 and B2 by days 44 and 25, respectively (Figure 2, red trajectories). A premature stop mutation (W307*) in *mutS* also rose to intermediate frequency (53%) in the P3 population by day 90. While we did not create an isogenic mutant of this mutator allele, we predict that it acts as other *mutS* mutations with a roughly 100-fold increase in mutation rate. The rise of *mutS* alleles caused these populations to diverge genetically from the others, as seen from PCA analysis of mutational frequency data (Figure 1B). A fourth mutator genotype evolved in the B3 population by a mutation in *mutL* (D467G) that caused a 16-fold increase in mutation rate and rose to 83% frequency by day 90 (Figure 2; Figure S5). Consequently, each of these mutator populations became enriched for transition and single-base indel mutations, signatures typical of defects in mismatch repair (Sniegowski et al. 1997; Shaver et al. 2002; Couce et al. 2017) (Table S2).

**Figure 2.**
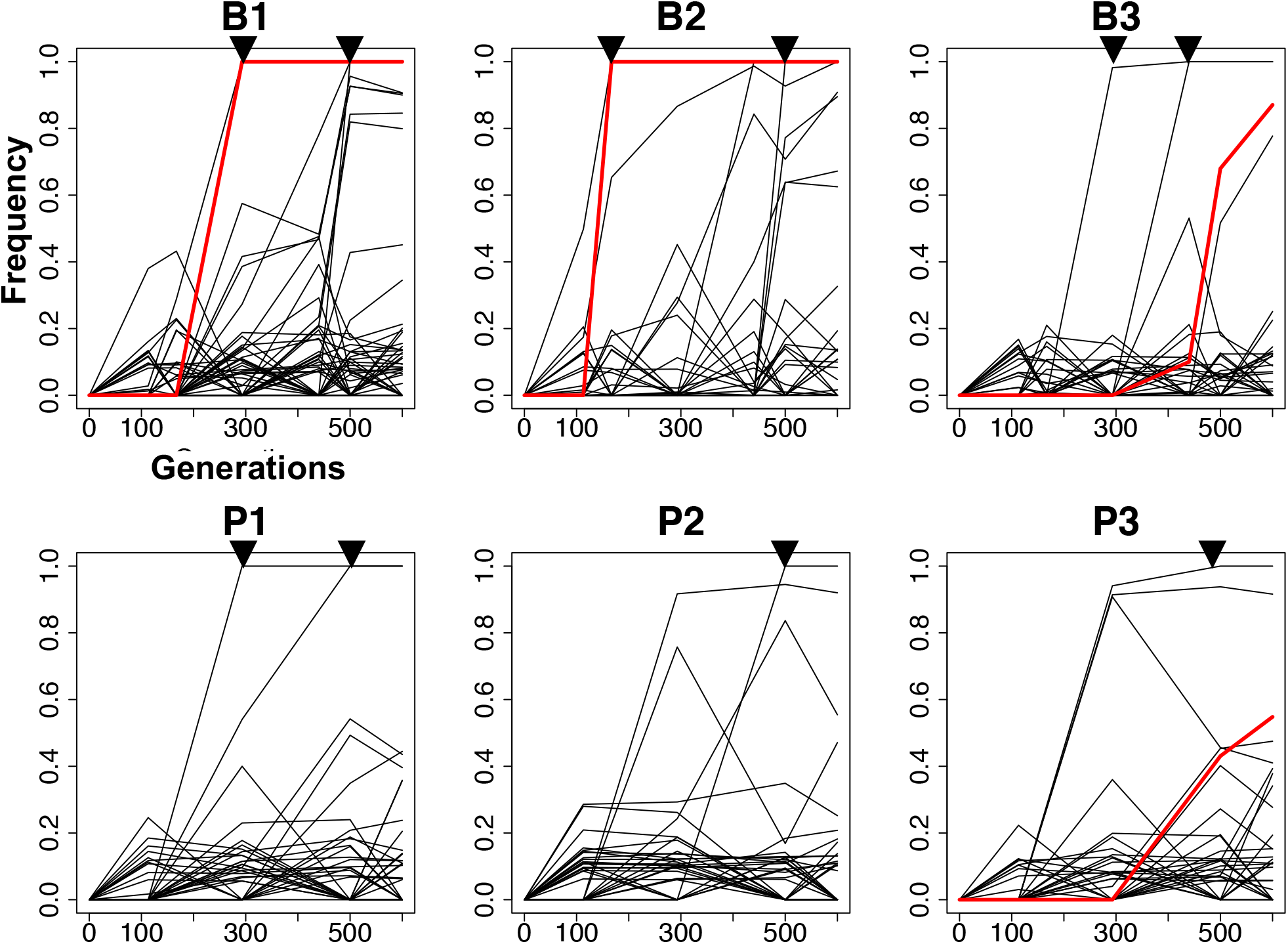
Evolutionary trajectories of inferred genotypes within six PA populations under biofilm (B) or planktonic (P) selection. Genotypes containing mutator alleles are colored red. Triangles indicate timing of the ten fixation events observed throughout the study.

### High diversity is a byproduct of clonal interference

While mutator alleles are not beneficial themselves (Sniegowski et al. 1997) and their dominant effect is to introduce neutral or deleterious mutations, they can facilitate the acquisition of combinations of beneficial mutations. A more beneficial genotype that has acquired several beneficial mutations may be able to outcompete less fit lineages with fewer mutations and overcome competition among adaptive lineages that slows their rate of increase. This dynamic is known as clonal interference or more generally as the Hill-Robertson effect (Birky and Walsh 1988; McVean and Charlesworth 2000; Comeron et al. 2008). The ability to acquire many beneficial mutations and escape clonal interference may be particularly advantageous when beneficial genetic variation is rare or, more likely in these populations, common (Smith and Haigh 1974; Buskirk et al. 2017).

As described above, hundreds of mutations rose to detectable frequencies in these populations within ∼200 generations and persisted. We used software recently developed by our lab to group mutations into genotypes on the basis of shared, nested frequency-trajectories (cdeitrick 2020). Remarkably, between 30 and 47 genotypes each composed of multiple mutations accumulated within each population throughout the experiment (Figure 2). The number of genotypes present at a given sampled timepoint continuously increased in biofilm populations, whereas the maximum diversity in planktonic populations occurred at day 44 (Supplemental Figure 4). Yet despite extensive variation at both nucleotide and genotype levels, only 10 fixation events were observed involving a total of 53 mutations. These selective sweeps involved genotypes with more mutations in biofilm populations than in planktonic lines (Table 1), and two sweeps were linked to mutator alleles in which 24 mutations fixed (Table S3). These results suggest that the biofilm environment selected larger mutational cohorts that could involve more differentiated genotypes to fix, whereas planktonic populations selected one or few mutations to fix. One cause of this pattern was the superiority of mutator genotypes containing more linked mutations, which were more common in biofilm lines. Combined, these findings suggest that the rapid rise of mutator alleles may have enabled escape from clonal interference, especially in biofilms.

To better understand the evolutionary processes within these populations, we visualized genotype frequencies over time by constructing Muller plots (Traverse et al. 2013; Scribner et al. 2019). These figures demonstrate, for example, how one genotype spreads by evolving secondary, nested, genotypes and outcompeting other preexisting genotypes. The most conspicuous genotype invasions were the mutator genotypes in populations B1 and B2 containing 11 or 13 mutations (Figure 3; mutator ancestry depicted in color and fixation events depicted by vertical lines). These figures also illustrate effects of particularly consequential mutations that overtake others or give rise to further diversity as well as genotypes that arise simultaneously and coexist for hundreds of generations. This latter dynamic is consistent with either clonal interference or genotypes adapting to inhabit discrete niches, possibilities we evaluate below.

**Figure 3.**
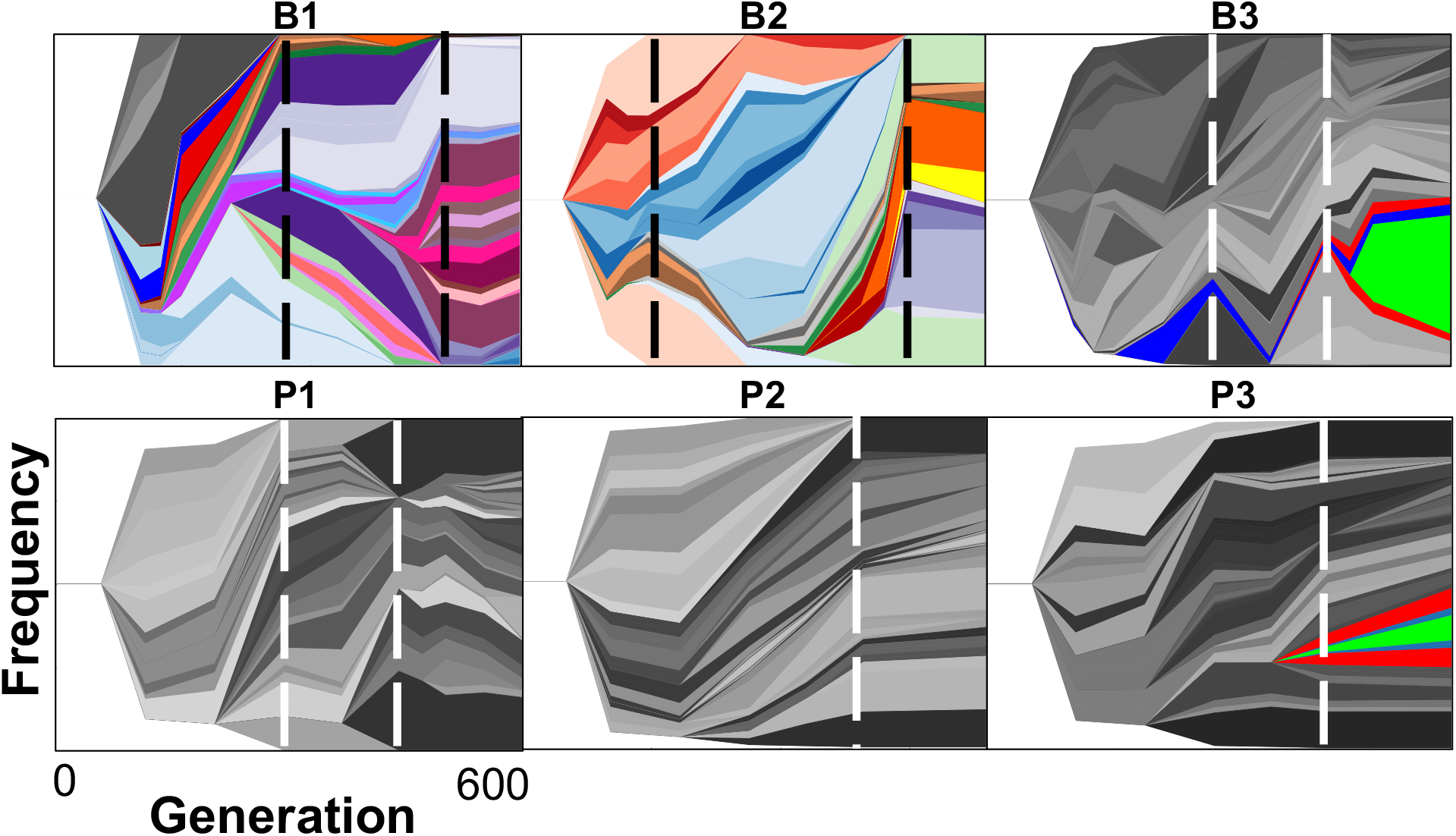
Genealogy and genotype frequencies over time. Each shade or color represents a different genotype and vertical area corresponds to genotype frequency, inferred by LOLIPop (cdeitrick 2020). Shades of grey indicate genotypes not on a mutator background, whereas colors indicate genotypes on a mutator background. Dashed lines indicate timing of fixation events predicted by genotype trajectories (Figure 2). Biofilm populations are on the top row and planktonic populations are on the bottom row.

To test the genotype prediction algorithm producing these Muller plots and to identify other mutations undetected due to their low frequencies or sequencing coverage, we sequenced 26 clones from population B1 and 10 from population P1. B1 clones contained 48-138 mutations per clone whereas clones from the P1 population only contained between 14-28 mutations, results consistent with population sequencing results. However, many of these mutations were previously undetected in the metagenomes, indicating much greater genetic variation at frequencies below our detection limits from population sequencing (Figure 4) (Good et al. 2017). The 26 B1 clones belonged to a total of nine competing genotypes at day 90 (Figure S6), whereas the 10 planktonic clones only belonged to two distinct genotypes (Figure S7). This finding confirms that the biofilm population maintained greater genetic diversity than the planktonic population at the end of the experiment.

**Figure 4.**
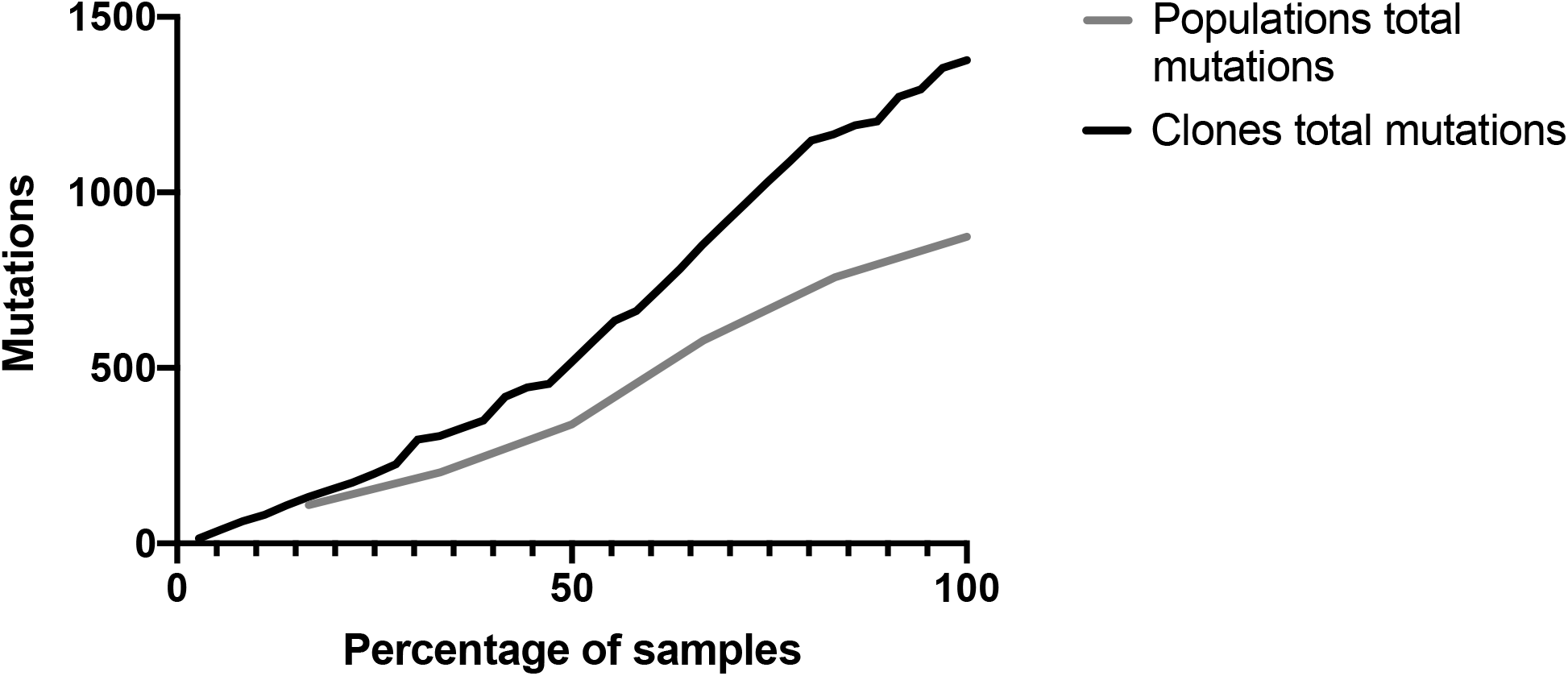
Clonal sequencing identifies more mutational variation than population level sequencing. Collector’s curve of total mutations identified by both population and clonal sequencing. To enable comparison results are plotted by the percentage of samples obtained (6 populations or 36 clones).

Another goal of sequencing these clones was to determine the genetic causes of the distinct colony morphologies that we previously showed to associate with distinct niches and phenotypes within the B1 biofilm population (Flynn et al. 2016). Each of the seven morphotypes (Figure 5a) represented a different long-lived genetic lineage (Figure 5b) that was separated by at least five mutations. This indicates the previous study that was based solely on colony morphotypes captured a good representation of the population diversity as it included seven of nine identified genotypes. We examined mutations unique to each genotype in an effort to identify those responsible for their unique phenotypes, which included the ability to form more productive biofilm communities in mixture (Table 2).

**Table 2.** Unique mutations found in seven clones from an experimentally evolved biofilm populations with distinct ecological roles(Flynn et al. 2016).

| Clone | Gene | PA01 ortholog | Pseudocap function |
| --- | --- | --- | --- |
| **A** | bkdA1 | bkdA1 |  |
|  | flgB | flgB | Motility & Attachment |
|  | oprC | oprC | Transport of small molecules |
|  | oprC | oprC | Transport of small molecules |
|  | PA14_06960 | <u>PA0534</u> | Putative enzymes |
|  | PA14_10830 | no PA01 ortholog | Transcriptional regulators |
|  | PA14_14530>/>PA14_14540 |  |  |
|  | gacS | gacS | Two-component regulatory systems |
|  | PA14_52760 | <u>PA0891</u> | Transport of small molecules |
|  | PA14_72320 | <u>PA5478</u> | Membrane proteins |
|  | ppc | ppc | Central intermediary metabolism |
|  | thrB>/<PA14_72520 |  |  |
|  | TrpB | TrpB | Amino acid biosynthesis and metabolism |
| **D** | oprH</>napE | oprH</>napE |  |
|  | PA14_04050 | PA0227 | Biosynthesis of cofactors, prosthetic groups and carriers |
|  | PA14_06030 | PA0461 | Fatty acid and phospholipid metabolism |
|  | PA14_10910 | PA4096 | Transport of small molecules |
|  | PA14_19450 | PA3453 | Hypothetical, unclassified, unknown |
|  | PA14_23670 | PA3127 | Antibiotic resistance and susceptibility |
|  | PA14_29260 | PA2696 | Transcriptional regulators |
|  | PA14_31470 | PA2557 | Fatty acid and phospholipid metabolism |
|  | PA14_48210 | PA1239 | Putative enzymes |
|  | PA14_55380</>PA14_55390 |  |  |
|  | magD | magD | Adaptation, Protection |
|  | PA14_66380 | PA5021 | Membrane proteins |
|  | pauR | pauR | Carbon compound catabolism |
|  | PA14_71750 | PA5437 | Transcriptional regulators |
|  | pvdJ | pvdJ | Adaptation, Protection |
| **H** | algU</>nadB | algU</>na<br>dB |  |
|  | dsbA2 | dsbA2 | Translation, post-translational<br>modification, degradation |
|  | gbuR | gbuR | Transcriptional regulators |
|  | PA14_19190 | PA3471 | Central intermediary metabolism |
|  | prkA | prkA |  |
|  | fliT | fliT | Motility & Attachment |
|  | gacS | gacS | Two-component regulatory systems |
|  | PA14_58960 | no PA01<br>ortholog | Hypothetical, unclassified, unknown |
|  | PA14_71750 | PA5437 | Transcriptional regulators |
|  | rpoB | rpoB | Transcription, RNA processing and<br>degradation |
| **M** | flhA | flhA | Motility & Attachment |
|  | PA14_07810 | PA0599 | Hypothetical, unclassified, unknown |
|  | PA14_13150 | PA3921 | Transcriptional regulators |
|  | PA14_43150</>PA14_43160 |  |  |
|  | pscQ | pscQ | Protein secretion/export apparatus |
| **O** | PA14_12160 | PA3992 | Putative enzymes |
|  | PA14_14390</>PA14_14400 |  |  |
|  | PA14_27610>/>PA14_27620 |  |  |
|  | PA14_27770 | PA2812 | Transport of small molecules |
|  | PA14_51510 | PA0988 | Hypothetical, unclassified, unknown |
|  | pstA | pstA | Transport of small molecules |
|  | spuF | spuF | Transport of small molecules |
|  | str | str | Antibiotic resistance and<br>susceptibility |
|  | trbL | trbL |  |
| **P** | lpxC | lpxC | Cell wall / LPS / capsule |
|  | mdoH | mdoH | Cell wall / LPS / capsule |
|  | PA14_03000 | PA0242 | Putative enzymes |
|  | PA14_13140>/>PA14_13150 |  |  |
|  | PA14_1340>/<PA14_13350 |  |  |
|  | PA14_18760 | PA3523 | Transport of small molecules |
|  | PA14_43150 | PA1652 | Membrane proteins |
|  | pilF | pilF | Motility & Attachment |
|  | purB | purB | Amino acid biosynthesis and metabolism |
|  | rep | rep | DNA replication, recombination, modification and repair |
| **V** | rsmE | rsmE | Translation, post-translational modification, degradation |
|  | pqiB | no PA01 ortholog | Hypothetical, unclassified, unknown |
|  | PA14_52250 | PA0929 | Two-component regulatory systems |
|  | rsmI | rsmI | Translation, post-translational modification, degradation |
|  | thrB>/<PA14_72520 |  |  |
|  | til5a>/>pIdA |  |  |

**Table 2.** Genes (n=40) with 3 or more evolved mutations. NS: nonsynonymous, S: synonymous, indel: insertion/deletion. Shading corresponds to frequency. Function from PseudoCap via pseudomonas.com.

| Gene | Total cases | Type |  |  | Biofilm | Planktonic | Highest frequency |  |  |  |  |  | Function |
| --- | --- | --- | --- | --- | --- | --- | --- | --- | --- | --- | --- | --- | --- |
|  |  | NS | S | Indel |  |  | B1 | B2 | B3 | P1 | P2 | P3 |  |
| argJ | 3 | 3 | 0 | 0 | 2 | 1 |  | 13.5 | 10.5 |  |  | 11.4 | Arginine biosynthesis |
| soxA | 3 | 2 | 1 | 0 | 1 | 2 |  |  | 10.6 |  | 14.7 | 11.4 | Carbon compound catabolism |
| aer2 | 5 | 3 | 2 | 0 | 3 | 2 |  | 9.6 | 10.9 | 10.2 |  | 14.2 | Chemotaxis |
| bdIA | 3 | 3 | 0 | 0 | 3 | 0 | 14.4 | 14.7 | 13.5 |  |  |  | Chemotaxis |
| cobG | 3 | 2 | 0 | 1 | 1 | 2 |  | 61.8 |  |  | 13.6 | 13.1 | Cofactor biosynthesis |
| mutS | 3 | 3 | 0 | 0 | 2 | 1 | 100 | 100 |  |  |  | 52.6 | DNA replication |
| napF | 3 | 3 | 0 | 0 | 2 | 1 | 13.8 |  | 10.9 |  |  | 10.7 | Energy metabolism |
| nqrE | 3 | 3 | 0 | 0 | 1 | 2 |  | 11.7 |  |  | 10.4 | 11.2 | Energy metabolism |
| nuoG | 3 | 3 | 0 | 0 | 1 | 2 | 12.9 |  |  | 12.1 | 13.9 |  | Energy metabolism |
| PA14_53110 | 7 | 7 | 0 | 0 | 2 | 5 |  | 14.8 | 12.8 | 12.9 | 14.4 | 12.1 | Putative enzymes |
| pslI | 7 | 7 | 0 | 0 | 3 | 4 | 16.7 | 16.4 | 14.6 | 16 | 14.7 |  | Putative enzymes |
| algF | 3 | 3 | 0 | 0 | 2 | 1 | 13.1 |  |  | 10.6 |  |  | Secreted Factors |
| fha1 | 4 | 4 | 0 | 0 | 3 | 1 | 27.6 |  | 25.1 | 35.8 |  |  | Secreted Factors |
| rpoB | 4 | 4 | 0 | 0 | 4 | 0 | 100 | 66 | 100 |  |  |  | Transcription |
| PA14_13150 | 3 | 3 | 0 | 0 | 3 | 0 | 46.3 | 17.8 | 100 |  |  |  | Transcriptional regulators |
| PA14_18200 | 6 | 6 | 0 | 0 | 5 | 1 | 22.3 | 22.1 |  | 12.8 |  |  | Transcriptional regulators |
| PA14_51840 | 3 | 3 | 0 | 0 | 0 | 3 |  |  |  |  | 12 | 16.6 | Transcriptional regulators |
| PA14_58510 | 5 | 3 | 2 | 0 | 4 | 1 | 23 | 15.7 |  | 15.5 |  |  | Transcriptional regulators |
| PA14_71750 | 3 | 3 | 0 | 0 | 3 | 0 | 93.9 | 89.7 |  |  |  |  | Transcriptional regulators |
| dsbA2 | 4 | 2 | 2 | 0 | 3 | 1 | 17.1 | 11.4 | 20.8 |  |  | 10.5 | Translation |
| rluB | 10 | 8 | 2 | 0 | 4 | 6 | 10.8 | 17.7 |  |  | 14.7 | 13.2 | Translation |
| nosD | 5 | 4 | 1 | 0 | 1 | 4 | 14.1 |  |  |  | 15.3 | 12.9 | Transporter |
| PA14_09300 | 3 | 3 | 0 | 0 | 2 | 1 | 16 |  | 15.2 | 18.6 |  |  | Transporter |
| PA14_22650 | 7 | 7 | 0 | 0 | 3 | 4 | 43.2 | 24 |  | 40 | 28 | 37.8 | Transporter |
| PA14_45060 | 6 | 6 | 0 | 0 | 2 | 4 | 24.4 | 19.1 |  | 19.5 | 14.1 | 13.6 | Transporter |
| PA14_46110 | 7 | 7 | 0 | 0 | 1 | 6 |  |  | 15.5 | 13.7 | 13.6 | 10.7 | Transporter |
| PA14_47900 | 3 | 3 | 0 | 0 | 0 | 3 |  |  |  |  |  | 19.9 | Transporter |
| pchF | 3 | 3 | 0 | 0 | 3 | 0 | 18.4 |  | 19 |  |  |  | Transporter |
| pitA | 4 | 2 | 0 | 2 | 2 | 3 |  | 91.1 |  | 100 | 75.8 | 100 | Transporter |
| pntB | 5 | 3 | 2 | 0 | 1 | 4 | 11.3 |  |  |  | 13.9 | 15.6 | Transporter |
| PA14_10770 | 3 | 2 | 1 | 0 | 1 | 2 |  | 100 |  | 12.2 |  | 26.3 | Two-component regulatory systems |
| PA14_32300 | 3 | 3 | 0 | 0 | 2 | 1 | 17.1 |  |  |  |  | 15.3 | Two-component regulatory systems |
| gacS | 3 | 3 | 0 | 0 | 3 | 0 | 85.2 | 92.2 |  |  |  |  | Two-component regulatory systems |
| PA14_01160 | 3 | 3 | 0 | 0 | 1 | 2 | 14.3 |  |  |  | 13.8 |  | Unknown |
| PA14_20510 | 6 | 6 | 0 | 0 | 3 | 3 | 9.6 | 11.5 |  | 11 | 11.9 | 11.1 | Unknown |
| PA14_31070 | 10 | 5 | 5 | 0 | 4 | 6 | 19.3 | 16.9 |  | 16.2 | 16.9 |  | Unknown |
| PA14_32830 | 6 | 6 | 0 | 0 | 2 | 4 |  | 13.8 | 12.7 | 12.1 | 13.5 | 12.1 | Unknown |
| PA14_54810 | 3 | 3 | 0 | 0 | 3 | 0 | 11.7 |  | 7.5 |  |  |  | Unknown |
| PA14_58070 | 6 | 6 | 0 | 0 | 5 | 1 | 13.6 | 17.5 | 12.9 |  | 20.8 |  | Unknown |
| PA14_69010 | 5 | 5 | 0 | 0 | 3 | 2 | 20.1 | 12.2 | 13 | 12.5 |  | 12.4 | Unknown |

**Figure 5.**
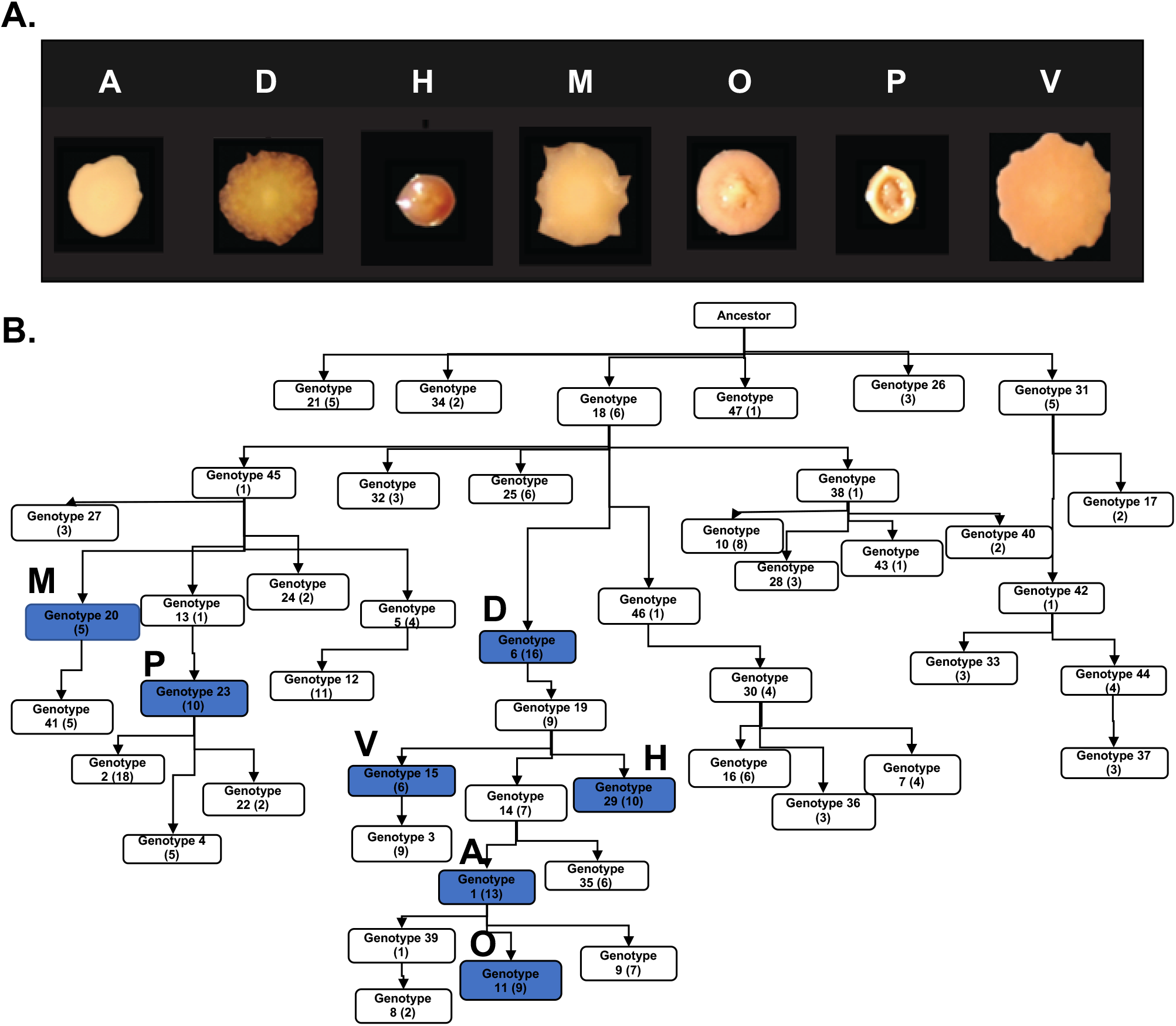
Clones representing different colony morphologies from the B1 population are all at different points on the evolutionary trajectory of the B1 population. (A) Seven colony morphologies were identified at day 90 and their phenotypes and ecological interactions were characterized previously(Flynn et al. 2016). (B) Ancestry was inferred using LOLIPop (cdeitrick 2020). Genotypes of named morphotypes are shown in blue.

For five morphotypes we can predict a genetic cause of the high biofilm formation and the related high c-di-GMP levels. Morphotypes A and H each independently acquired mutations in *gacS*, a sensor kinase known to be associated with high biofilm production(Davies et al. 2007), and which will be discussed more below. The additional three evolved mutations in separate genes that have all been linked to *Pseudomonas* biofilm production previously: mutant D acquired a mutation in the sensor histidine kinase *rcsC* (Nicastro et al. 2009), M acquired a mutation in the type III secretion system gene *pscQ* (Wagner et al. 2003), and V acquired a mutation in the two component response regulator PA14_52250 (Francis et al. 2017; Badal et al. 2020) (Table 2). While these are not the only mutations that distinguish the morphotypes from one another, these are leading candidates that could produce their varied biofilm phenotypes.

### Targets of selection

Evolving large populations (>10^8^) over a few hundred generations results in higher impacts of selection relative to the effect of drift, even when mutation rates increase. This allows us to predict that genotypes that rise to a detectable frequency in an evolve-and-resequence experiment like this one are fitter than their ancestor. Further, parallel, nonsynonymous mutations affecting the same gene provides strong evidence of an adaptation (Cooper 2018). Across all six populations, 53 mutations reach 100% frequency, as part of the 10 genotype sweeps (Table 1), and each affected a unique gene except for two each in *mutS, pitA*, and *rpoB* (Table S3). However, the population dynamics depicted in Figure 3 and the clonal genomes summarized in Figure 5 demonstrate substantial genetic variation that did not fix. We suspected a more appropriate screen for parallelism might involve a lower frequency threshold than 100%. At 80% frequency, eight genes were mutated more than once: *lysC, mutS*, the intergenic region *fabI/ppiD*, PA14_23670, *pitA*, PA14_69980, PA14_71750, and *rpoB*, and all six populations are represented. Mutations affecting these genes were either small indels or nonsynonymous substitutions predicted to alter or eliminate function. Because these genes were mutated in both planktonic and biofilm environments, they likely enable adaptation to the growth medium, temperature, and dilution timing. However, these represent only 17 of the 116 mutations that reach 80% frequency, a remarkably flat distribution of high-frequency genotypes that suggests many loci may provide a benefit under these conditions.

We next analyzed the complete set of parallel mutations found at all frequencies for those passing a statistical threshold of expected random co-occurrence and identified 153 genes or intergenic regions. Forty genes were mutated three or more times, more than would be expected by chance, accounting for 179 mutations (21.1% of all mutations; Table 3). Most mutations are nonsynonymous, as expected from positive selection, and many suggest possible adaptive phenotypes.

The gene with the most independent mutations, *rluB* (PA14_23110) with 10 (8 nonsynonymous; Table 3, Table S3) found in four populations at frequencies of 12.1-17.7%, is difficult to understand as an adaptation. *RluB* encodes a 23s rRNA pseudouridine synthase responsible for modifying this rRNA at position 2605 during maturation (Kaczanowska and Rydén-Aulin 2007). This gene was mutated in both biofilm and planktonic populations, indicating it is not specific to lifestyle, but all mutations clustered in residues 313 and 323 found in the disordered C-terminus of this protein. In addition, one mutation in *rluC*, another pseudouridine synthase, was detected. The literature remains vague about the specific phenotypic roles of these 23S rRNA modifications and effects of disrupting these genes are difficult to detect *in vitro*, but a recent study implicated these modifications as important for growth under anoxia (Krishnan and Flower 2008; Basta et al. 2017).

Fortunately, a wealth of literature has identified genes involved in the transition from planktonic to biofilm growth in *P. aeruginosa* (Boles et al. 2004; Eichner et al. 2014). These genes are often well annotated at http://pseudomonas.com for the PA14 genome or in related strains (Winsor et al. 2016). For example, five mutations in PA14_02220, encoding the aerotaxis transducer Aer2, were selected in two biofilm and one planktonic population, and may alter the response to low oxygen that is common in biofilms (Sawai et al. 2012). In addition, three independent V334G mutations were selected in each biofilm population in PA14_46030, encoding the ortholog of <u>b</u>iofilm <u>d</u>ispersion locus BdlA (Morgan et al. 2006). This cytoplasmic protein contains two sensory PAS domains and a chemoreceptor domain that has been shown to sense and mediate responses to oxygen levels (Petrova and Sauer 2016). Two-component regulatory systems are also commonly implicated in this lifestyle transition, and in agreement with this model, 19 mutations in 14 genes encoding sensor kinases, response regulators, or hybrid complexes were identified (Table 3, Table S3). Likewise, nine different nonsynonymous or indel mutations affecting motility and attachment (e.g. *flgB/G, fliC/H/M, pilB/pilQ*, and *cupB3*) were selected in both biofilm and planktonic populations, indicating advantages of disrupting these processes in both lifestyles.

More surprising, the most common functional classification of mutated genes was transporters. Nine transporters were mutated repeatedly including: *pitA, pntB, nosD, pchF*, PA14_09300, PA14_22650, PA14_45060, PA14_46110, and PA14_47900, and account for 24% (43) cases of parallelism (Table3, Table S3). These transporters include genes with known function including inorganic phosphate transporter *pitA* and copper transporter *nosD*, as well as those with unknown substrates including PA14_46110 and PA14_45060. Of all transporters identified, *pitA* was the only one to reach frequencies above 50% in any population. This led us to identify an additional stressor acting on these populations as a byproduct of growth on arginine as a sole carbon source, which is an alkaline pH (pH = 9.2 at 24 hr) caused by the release of ammonia through deamination (Ha et al. 2014). It remains unclear how these mutated transporters produce adaptations, but they could be acting to balance pH or compensate for redox imbalances or changes in proton motive force. In addition, 18 mutations affected genes involved in various metabolic processes, including *argJ*, a key component of arginine biosynthesis expected to be no longer required due to abundant arginine in the growth media, *soxA*, encoding sarcosine oxidase (Willsey and Wargo 2016), and *cobG*, involved in the aerobic pathway of cobalmin (Vitamin B12) biosynthesis (Blanche et al. 1992). Further, three mutations are observed in each of *napF, nqrE*, and *nuoG* potentially altering the energy conserving respiratory NADH dehydrogenase chain (Hoboth et al. 2009). We suspect that many of these nonsynonymous or indel mutations are partial loss of function mutations in metabolic pathways that are extraneous in the M63+arginine medium, though an exact mechanism is unknown.

There are four notable cases of mutated genes that are lifestyle specific. Four genes (*rpoB, gacS*, PA14_71750, and PA14_13150) are only mutated in biofilm populations, and in all cases, mutations rise to greater than 90% in at least one population (Table 3, Table S3). Mutations in *rpoB* and other RNA polymerase genes (*rpoACD*) have been frequently identified as adaptations during evolution experiments (O’Sullivan et al. 2005; Xiao et al. 2017). One explanation in *Escherichia coli* suggests they are adaptive for growth in minimal media by redistributing RNA polymerase from small RNA promoters (i.e. rRNA) to rate-limited promoters required for anabolic processes from a limited set of carbon sources, such as arginine in the case of this study (Conrad et al. 2010). GacS is a sensor/regulator hybrid known to govern a broad range of traits involved in virulence, secondary metabolism and biofilm formation through the regulation of small rRNA’s (Gellatly et al. 2018). Further, *gacS* has been shown to be involved in the switch from a hyperadherent small colony phenotype back to a wild type phenotype indicating that loss of GacS function is beneficial in biofilms(Davies et al. 2007). The additional two biofilm specific loci, PA14_13150 and PA14_71750, are less understood transcriptional regulators, however PA14_71750 has been previously associated with biofilm adaptation in *Pseudomonas* (Konikkat et al. 2020).

The rarity of fixation events and the high incidence of genes with parallel mutations throughout the experiment provides further evidence of competition between adaptive lineages in a population, or clonal interference (Lang et al. 2013). Given a set of genes in which mutations are adaptive, we would expect each competing genotype to contain different combinations of these mutations, with the most beneficial of them repeatedly evolving on different genetic backgrounds. As predicted, we identified 55 cases of within-population gene level parallelism (4 – 16 cases per population), with most (87%) predicted to be on different genotypes. This result also demonstrates the polygenic capacity for adaptation by *P. aeruginosa* growing in minimal arginine medium.

### Selected genotypes are adapted to both growth conditions and lifestyle

Most cases of gene-level parallelism were shared between biofilm and planktonic conditions and indicate adaptations to the common growth media. However, minority variants could include environment specific adaptations. For example, our bead transfer model simulates the complete life cycle of the biofilm – attachment, assembly, dispersal, and reattachment – each of which could select for discrete phenotypes. To distinguish genotypes adapted to either biofilm or planktonic growth, we grew 90-day samples of evolved populations B1, B2, P1, and P2 for two days in each of these conditions independently (Figure 6). We hypothesized that these treatments would enrich genotypes adapted to either condition that we could resolve by resequencing treated populations and correlating their relative frequencies. Mutation frequencies that are higher in one environment, could represent genotypes that are adapted to biofilm or planktonic growth.

**Figure 6.**
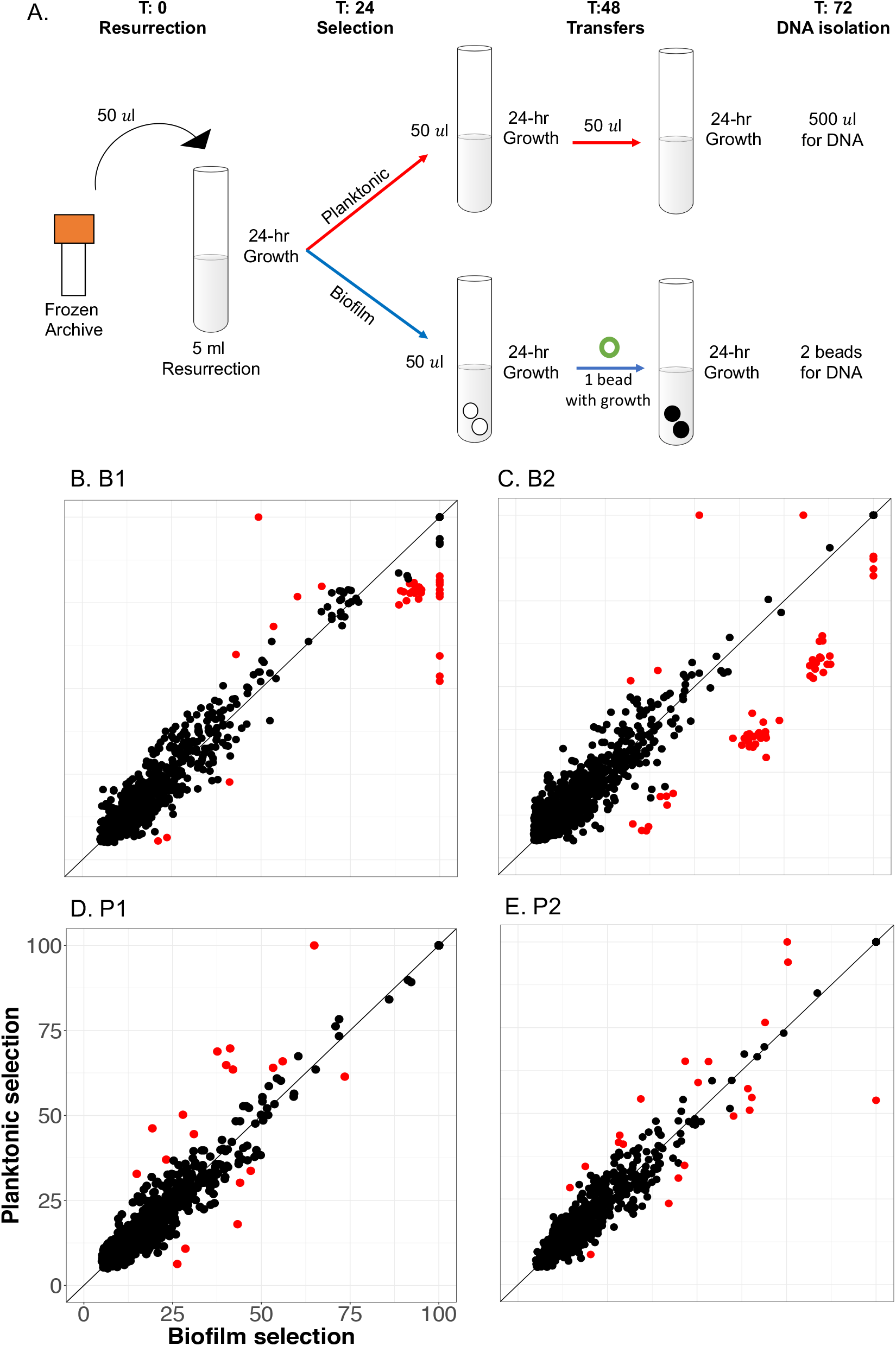
Ecological signals from WGPS data after two days of selection enriches mutations adaptive in both planktonic and biofilm conditions. A. Methods for isolating DNA to identify genotype by environment interactions. 90-day populations were resurrected from frozen archives in planktonic environments for 24 hours. Each population was then split and subjected to two days of either planktonic selection, via 1:100 daily transfers, or biofilm selection, via the bead model. After the second day of selection, 0.5 ml of planktonic culture and two biofilm beads were used to obtain the planktonic and biofilm DNA samples, respectively (see methods). B-E. Correlation of mutational frequencies when resurrected population DNA is isolated after two days of planktonic selection versus biofilm selection. B: B1 population, C: B2 population, D: P1 population, and E: P2 population. Mutations that are significantly enriched via Cook’s distance (see methods) are represented in red and other mutations in black.

In all four populations tested we found significant enrichment in both biofilm and planktonic environments (Figure 6 b-e). As expected, the two biofilm populations tested had more biofilm enriched mutations than planktonic enriched (37 and 55 biofilm enriched compared to 4 and 2 planktonic enriched), though planktonic populations had roughly equal numbers of biofilm and planktonic enriched mutations (8 and 10 biofilm enriched versus 10 and 11 planktonic enriched; Table S4). Upon closer examination, the B2 population appears to contain a set of 36 mutations belonging to 4 genotypes enriched for biofilm fitness (Table S4). These genotypes also rose in frequency between days 75 to 90 in the evolution environment and appear to be generally adaptive. However, these experiments did not generally cause large (>10%) shifts in genotype frequency, which suggests that they correspond to adaptations to the overall experimental conditions and not just these growth phases.

## Discussion

We used experimental evolution to examine how the opportunistic pathogen *Pseudomonas aeruginosa* (PA) adapts in laboratory culture in a medium known to promote biofilm formation (Ha et al. 2014), comparing the results of propagation by simple serial dilution with a model simulating the biofilm life cycle (Poltak and Cooper 2011). After 90 propagations (∼600 generations) we observe more mutations per population, fewer fixation events, and earlier prevalence of mutator genotypes than has been observed in other evolution experiments. These findings are consistent with two potentially overlapping processes: 1) the existence of many coexisting niches, representing differing metabolic or ecological strategies within the laboratory environment that select for a variety of specialized genotypes or 2) high clonal interference, involving many genotypes with roughly the same fitness competing against one another in the same niche. The repeated evolution of mutations in the same gene in the same population provides the clearest evidence of this clonal interference, because such mutants are likely to share similar phenotypes. For example, within the B1 population mutations in *gacS* evolve independently in two genotypes (Table S3). We confirmed this finding through clonal sequencing, finding clones H and A to each contain a *gacS* mutation in their distinguishing genotypes (Table 2), and further, these clones were previously shown to compete for a common niche (Flynn et al. 2016). While clonal interference likely contributes to the high genetic variation observed in these populations, we also predict that multiple niches exist within each population as our previous study indicated (Flynn et al. 2016). As further evidence, most cases of gene-level parallel mutations reached only 20-30% frequency within their respective populations (Table 3). We hypothesize that this results from multiple coexisting ecotypes with different adaptations within each population, a phenomenon that has been shown in previous evolution studies, particularly with biofilms (Traverse et al. 2013; Buskirk et al. 2017; Good et al. 2017).

To identify the traits selected by growth in a minimal medium containing arginine as sole energy source, we focused on the 40 loci mutated most frequently (Table 2). This medium was selected because it had previously been shown to induce high biofilm production by *Pseudomonas* (Ha et al. 2014) more specifically by increasing intracellular levels of the secondary messenger molecule c-di-GMP through the two diguanylate cyclase proteins RoeA and SadC. Further, arginine was the only amino acid to completely repress swarming motility, indicating this carbon source could be a cue to enter an attached lifestyle (Bernier et al. 2011). As we were interested in seeing how *P. aeruginosa* continues to evolve under conditions where it is already proficient for biofilm, this media choice was suitable. However, it is a naturally stressful environment because arginine catabolism by arginine deiminase produces ammonia as a byproduct and hence an alkaline environment (Cunin et al. 1986; Lu 2006). These conditions are opposed to previous evolution experiments with *P. aeruginosa* in which media with multiple carbon sources are used to mimic an infection, and the medium often becomes acidic (Bernier et al. 2011; Wong et al. 2012; Ha et al. 2014; Scribner et al. 2019). The exact effects of arginine as the sole carbon and nitrogen source on the metabolic evolution of *P. aeruginosa* are unclear and the topic of a future study, however it is likely that arginine as a sole carbon and nitrogen source produced strong selection and may have been the dominant pressure.

The remarkable, unexpected consequence of this experimental design was a lack of convergent evolution of a small number of genes in replicate populations. Rather, dozens or even hundreds of mutations in nearly as many genes appear to have been selected, suggesting polygenic adaptation involving many metabolic and regulatory systems (Figure 3). Such polygenic selection in large populations increases the probability of clonal interference, wherein each lineage acquires its own combination of adaptations, but any given genotype would struggle to become dominant. Consequently, fixation events were rare, with only ten across all six replicate populations over 600 generations. Further, one of the most commonly mutated pathways was DNA mismatch repair, which increased mutation rates of linked genotypes in the four populations in which they evolved. We hypothesize these mutator genotypes provided the means to escape clonal interference (Good et al. 2014) because they had a higher probability of producing multiple, linked beneficial mutations that could outcompete other genotypes, a phenomenon that has previously been associated with the introduction of alternative forms of structure (Raynes et al. 2019).

Our previous study of these populations predicted that the different colony morphotypes represented lineages that had evolved to occupy distinct, interacting ecological niches within a biofilm population (Flynn et al. 2016). This genomic study confirms that these genotypes indeed represent different long-lived lineages within the B1 population. Each clone was found to possess numerous evolved mutations that are candidates for the observed phenotypic variation reported previously, e.g. fitness versus the ancestral PA14 strain, levels of cyclic-di-GMP, and motility (Flynn et al. 2016). However, considerably more variation was detected by population sequencing than these lineages could explain, and likewise, sequencing additional clones resulted in an ever-increasing census of new mutations (Figure 4). These results reinforce the polygenic model of adaptation to the biofilm-inducing arginine medium and the diversity of genotypes that can generally improve growth and survival or inhabit the niches illustrated in our prior study (Flynn et al. 2016; Barghi et al. 2020). More generally, this work has implications for understanding processes underpinning the evolution and maintenance of high levels of genetic diversity, including how different nutrient sources influence this process. Although differences in the mode of growth have been shown to influence genetic diversity (O’Toole et al. 2000; Boles et al. 2004; Resch et al. 2005; Booth et al. 2011), the metabolic environment may be equally or even more important by exposing varied fractions of the genome to selection. Further, this study provides a concrete example of how microbial populations evolving in a new environment can adapt and diversify without the periodic losses of genetic variation caused by selective sweeps. This dynamic was demonstrated in the Long-Term Evolution Experiment with *E. coli* after many thousands of generations had passed and the rate of adaptation had slowed (Couce et al. 2017; Good et al. 2017). Yet here, populations amassed extreme fitness gains over only hundreds of generations while maintaining many lineages, suggesting that the *P. aeruginosa* genome encodes vast potential to meet new environmental challenges.

## Materials and methods

### Experimental evolution

Replicate populations were experimentally evolved as previously reported (Flynn et al, 2016). Briefly, the ancestral *Pseudomonas aeruginosa* (PA14; NC_008463) strain was reconstituted from a freezer stock in Luria-Bertani broth (LB; 1.0% w/v tryptone, 0.5% w/v yeast extract, 1.0% w/v NaCl). Six replicate populations were started with 1:100 dilutions of an overnight growth into 5 mL M63 media (15 mM (NH_4_)_2_SO_4_, 22 mM KH_2_PO_4_, 40 mM K_2_HPO_4_, 40 mM galactose, 1 mM MgSO_4_, 25 μM FeCl_2_, and 0.4% w/v L-arginine(Bernier et al. 2011)). Three populations were propagated under liquid grown, planktonic, selection (P1, P2, P3), while three populations were propagated, concurrently, under constant biofilm selection (Poltak and Cooper 2011; Traverse et al. 2013; Flynn et al. 2016) (B1, B2, B3). Planktonic selection consisted of daily 1:100 liquid dilutions into 5 mL M63 media, while biofilm selection consisted of the transfer of one 7 mm polystyrene bead every 24 hours to a new tube of 5mL M63 media containing one clean bead. This biofilm selection method requires the entire biofilm lifecycle of dispersal, attachment, and growth between every transfer. All six populations were propagated for 90 days (∼600 generations) in 18×150 mm test tubes at 37 °C on a roller drum at 30 rpm.

Archives were made for all populations at days 17, 25, 33, 45, 66, 75, and 90 in 8% dimethyl sulfoxide (DMSO) at -80 °C. Planktonic populations were archived by freezing a 1 ml aliquot of the 24-hour culture, whereas biofilm populations were archived by sonicating 48 hr beads in 1 ml PBS, and then freezing the PBS with 8% DMSO.

### Phenotypic characterization of evolved populations

All populations were revived from freezer stocks for characterization by adding 50 µl of freezer stock to a culture of 4:1 M63 media to LB media, with trace elements to recreate previous water mineral content (110.764 g/L CaCl_2_, 0.824 g/L MnSO_4_, 33.48 g/L KBr, 132.737 g/L Na_2_SiO_3_, 123.485 ZnSO_4_, 0.178 g/L CoCl_2_ *6H_2_O, 1.458 g/L CuSO_4_). This media combination was used to minimize selection effects of growing in a fully complex medium, while providing enough nutrients for the frozen cells to revive. Resurrected populations were grown for 24 hr at 37 °C on a roller drum before being vortexed for 10 seconds, twice, to ensure biofilm growth was disrupted from the sides of the glass tube.

Growth curves were measured for resurrected populations in 96 well plates, at 37 °C. All growth curves were started at an OD_600_ of 0.01 and growth was measured every 10 min following 9 min of shaking for 24 hr in M63 media with trace elements. Maximum growth rate was determined for the average of all replicates, by using the following equation:

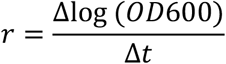

Census population size was calculated by diluting resurrected populations 1:100 in 5 mL M63 media with trace elements. One 7 mm polystyrene bead was added to each biofilm culture at time zero. After 24 hr growth, all populations were vortexed for 30 seconds and then diluted and plated in triplicate on tryptic soy agar and grown at 37 °C for 48 hrs. Average colony counts of the three technical replicates were reported for each population. Additionally, populations plated for census population size were also used for pH measurements as they were complete replications of experimental conditions. Average pH of triplicate readings are reported.

Biofilm assays were performed on resurrected populations diluted to an OD_600_ of 0.01 in M63 media and grown in a 96 well plate under static conditions for 24 hr at 37 °C. Media was discarded and the plate was washed twice with deionized H_2_O, which was also discarded. Wells were then stained with the addition of 250 μl 0.1% crystal violet solution for 15 minutes. Plates were rinsed to remove excess dye and the plate was allowed to dry for 24 hours. A de-stain solution (95% ethanol, 4.95% dH_2_O, and 0.05% Triton X-100 (fisher bioreagents)) was added to wells (250 μl per well), and, after a 15-minute incubation, was transferred to a new 96 well plate. Crystal violet absorbance readings were measured at 590 nm. Reported values are the average of 21 replicates, done across three plates (7 technical replicates per plate). Data were analyzed with a one-way ANOVA with post hoc Tukey test.

Motility assays were performed as described previously (Ha, et al 2014). Briefly, pipet tips were dipped into resurrected populations and stabbed into motility agar (0.3% agar, 6 g/L Na_2_HPO_4_, 3 g/L KH_2_PO_4_, 0.5 g/L NaCl, 0.2% glucose, 0.5% casamino acids, 1 mM MgSO_4_). Increased or decreased motility was determined for all day 90 evolved populations as the diameter of bacterial growth after 24 hr growth at 37 °C compared to the clonal ancestor. Reported values are the average of 5 replicates. Results were analyzed by one-way ANOVA with post hoc Tukey test.

### Mutation rate estimate

Isogenic mutants of biofilm evolved mutator alleles *mutS* T112P and *mutL* D467G were used as reported previously(Flynn et al. 2016). Briefly, the two mutator strains and the PA14 ancestor were revived from freezer stocks by streaking on ½ trypic soy agar. After 24 hrs growth individual colonies were used to start 30 replicate 5-ml cultures for each strain. After 24 hrs growth at 37° F on a roller drum, populations were diluted and plated on ½ tryptic soy agar both with and without 1mg/ml Ciprofloxacin as a selective agent. Colonies were counted after a 24 hour incubation period on both the antibiotic and non-antibiotic plate. We then calculated the fold change in mutation rate using the maximum likelihood method of Gerrish (2008). We performed measurements for all strains simultaneously to minimize variation. Reported values are the average fold change of all replicates.

### Genomic sequencing of evolved metagenomes

DNA was isolated from biofilm populations at 113, 167, 293, 440, 500, and 600 generations (17, 25, 44, 66, 75, and 90 days) and from planktonic populations at 113, 293, 500, and 600 generations (17, 44, 66, and 90 days). Culture media for DNA isolation was composed of 4:1 M63 to LB media. DNA was isolated from 1 mL resurrected populations using Qiagen’s DNeasy Blood & Tissue Kit. Library construction was done using the Illumina Nextera kit as described previously (Baym 2015). Libraries were sequenced on Illumina’s NextSeq 500 platform by the Microbial Genome Sequencing Center at the University of Pittsburgh. Between 13,545,344 and 46,890,487 reads were obtained for each sample, resulting in coverage of 265x - 860x per sample.

### Determining mutational frequencies from sequencing data

Initial metagenomics sequencing efforts of the six populations resulted in 121.2 gbps of 2×151 bp sequencing reads, that we trimmed and quality filtered with trimmomatic v0.36 using default parameters(Bolger et al. 2014). The breseq software package v0.31.0 was used to align the filtered reads to the reference PA14 genome (NCBI’s RefSeq database: NC_008463.1) and make polymorphism calls(Barrick et al. 2014). All populations were run using the polymorphism mode of breseq with a sliding window of 5 bp and a quality score cutoff of 10. This analysis was done for every population and every time point sequenced. The genome of the ancestral strain differed from the reference by 435 mutations, which were removed from all evolved population samples before any downstream analysis to remove mutations that did not evolve over the course of the evolution.

All 12,250 resulting mutation calls were filtered in R (v3.5.1, ^5^) to require a mutation to be called by at least 3 reads on each strand (positive and negative). Finally, mutations were filtered to remove regions of high polymorphism. Due to the abundance of repetitive regions in the PA14 genome the false positive rate is high. We therefore examined each call using the alignments produced by breseq and excluded mutations with high sequence variation within the 15 bases both upstream and downstream of the called mutation. Mutations were also filtered for known regions of misalignment such as repeat regions before consolidation into time-series tables containing all real mutated loci for each evolved population (Table S1). The resulting 874 mutations were then used as input in the software package LOLIPop Version 0.8.1. using – similarity-cutoff value of 0.1 for P1, P2, P3, and B1 and a --similarity-cutoff of 0.2 for populations B2 and B3. Increased similarity cutoffs increased the amount of mutations that were grouped into a genotype. Types of mutations were broken down by both nucleotide and amino acid mutation for all evolved populations in R. All scripts for the R filtering and analysis are available on github at: https://github.com/KatrinaHarris23/PALTEanalysis

### Calculating dN/dS ratios

To determine the dN/dS ratios, first the dN/dS ratio for each of the 64 codons was calculated. The codon usage for the ancestral, PA14, genome was obtained from (http://www.kazusa.or.jp/codon/). The two resulting matrices were multiplied to obtain a neutral ratio of nonsynonymous to synonymous substitutions (dN/dS ratio) of 2.96 for the ancestral UCBPP-PA14 genome (script for calculating this can be found on github here https://github.com/KatrinaHarris23/PALTEanalysis/blob/master/dnds_calculator). All reported dN/dS ratios are standardized according to this number with the final number reported being a result of the following equation:

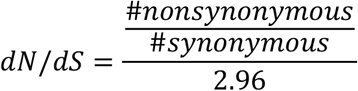

### Experimentally determining ecologically enriched mutations

To determine if mutations were specific to the biofilm lifestyle steps of swimming or attaching, we isolated DNA from evolved populations after additional strong selection in two distinct environments (Figure 5). Populations were resurrected from day 90 archives for the B1, B2, P1, and P2 populations by using 50 ul of the frozen stock into 5 ml resurrection media. After 24 hr growth at 37°C each culture was used to inoculate 2 identical 5 ml M63 arginine media cultures. Two 8mm polystyrene beads were added to one replicate, creating movable biofilm, before both inoculated cultures were placed at 37°C. One transfer was performed for all populations at 24 hr. Swimming was selected for by transferring liquid phase growth form replicates with no beads to a new tube with 5 ml M63 arginine media. Additionally, one bead from replicates with beads was rinsed with PBS and subsequently transferred to a fresh tube of 5 ml M63 media with two differently colored beads. After a final 24 hr incubation at 37°C “planktonically enriched” DNA samples were isolated from 0.5 ml liquid phase from replicates without beads. Growth from the two colored beads, first rinsed in PBS, then sonicated in 1 ml PBS was used in total for our “biofilm enriched” DNA sample. DNA was isolated using the Quiagen DNeasy Blood and Tissue kit.

Between 328 - 410x coverage was obtained for all eight samples. Mutations were filtered by removing all ancestral mutations, and only mutations detected by both biofilm and planktonic samples for a given population were considered. This sequencing effort resulted in 979, 1021, 879, and 876 mutations in the B1, B2, P1 and P2 populations respectively. Relative frequencies of mutations in the two environments were plotted in R using the ggplot2 package (Figure S8; code used in this analysis is found on github https://github.com/KatrinaHarris23/PALTEanalysis/blob/master/201020_ecological_interactions. <u>R</u>).

### Clonal sequencing

To acquire clonal DNA, populations were resurrected in 4ml M63 arginine media with 1ml LB and grown for 24 hr at 37 on a roller drum. Populations were then plated on ½ Tsoy plates down to the -6 dilution to isolate individual colonies. After 24 hr growth and 37 followed by 48 hr room temperature growth clones were selected ensuring that all visible colony morphologies were sampled. Colony growth was picked up and used as inoculum for 5 ml LB cultures. After 24 hr at 37 on a roller drum 1 ml of culture was used for archiving (9% DMSO) and 0.5 ml was used for DNA isolation using the DNeasy Blood and tissue kit. Named clones that were reported previously(Flynn et al. 2016) were already archived and archives were used for resurrection directly into 5 ml LB.

DNA sequencing was performed as with population sequencing, but only requiring 30x coverage per sample. All reads were trimmed with trimmomatic, as populations were, and variant calling was performed using Breseq software default settings. As clonal samples are only looking for mutations that fix in each sample, filtering was only performed to remove ancestral mutations before analysis.

### Statistical testing

Alpha diversity was calculated for all detected mutations in each population at each time point. We used the “shannon”, “simpson” and “invsimpson”, modes of the diversity function in R. Populations were grouped by growth environment, biofilm (n = 3) and planktonic (n = 3), and by mutator allele presence (n = 4) or absence (n = 2). T-tests were performed in Prism (version 8, www.graphpad.com, Figure S4).

To perform the Fisher’s exact test, a table of loci observed mutated 2 or more times was generated from the final mutation calls. Locus length was determined using http://pseudomonas.com as either gene length or the distance between surrounding genes for intergenic regions. Fisher’s exact test was performed using this compiled list: the numbers of mutations per loci, the total number of mutations observed (624), and the PA14 genome length (6537648). Testing was performed using the fisher.test() function in R and only loci with p-values lower than 0.05 were used for parallelism analysis. 153 loci were tested for parallelism and then the number of false positives were reduced using the Benjamini-Hochberg correction with a false discovery rate of 5%. The equation used was (i/m)Q where i is the p-value rank, m is the total number of tests, and Q is the false discovery rate.

Finally, ecological enrichment significance was calculated in R by using Cook’s distance. First, a linear regression was calculated for each population using planktonic versus biofilm frequencies using the lm() function. Cook’s distance was then calculated for each point, using cooks.distance() function. A mutation was considered significantly enriched if its distance was four times the mean distance or more.

### Principal component analysis (PCA)

All PCA calculations and plotting were performed in R. Principal components were computed from the biofilm production, swimming motility, and maximum growth rate (Vmax) data, and from the mutational frequency data at each sampled day using the prcomp() function. Plotting was done using the autoplot() function that is part of the ggfortify package.

## Acknowledgements

This work was supported by NASA CAN-7 NNA15BB04A and a Research Development Pilot award from the Cystic Fibrosis Foundation to VSC. We thank Dan Snyder and Ann Donnelly for critical feedback and Christopher Deitrick for support in implementing the lolipop package.

## Notes

https://github.com/KatrinaHarris23/PALTEanalysis

### Competing Interest Statement

The authors have declared no competing interest.

