## Supplementary Materials for "Polygenic adaptation, clonal interference, and the evolution of mutators in experimental *Pseudomonas aeruginosa* populations"

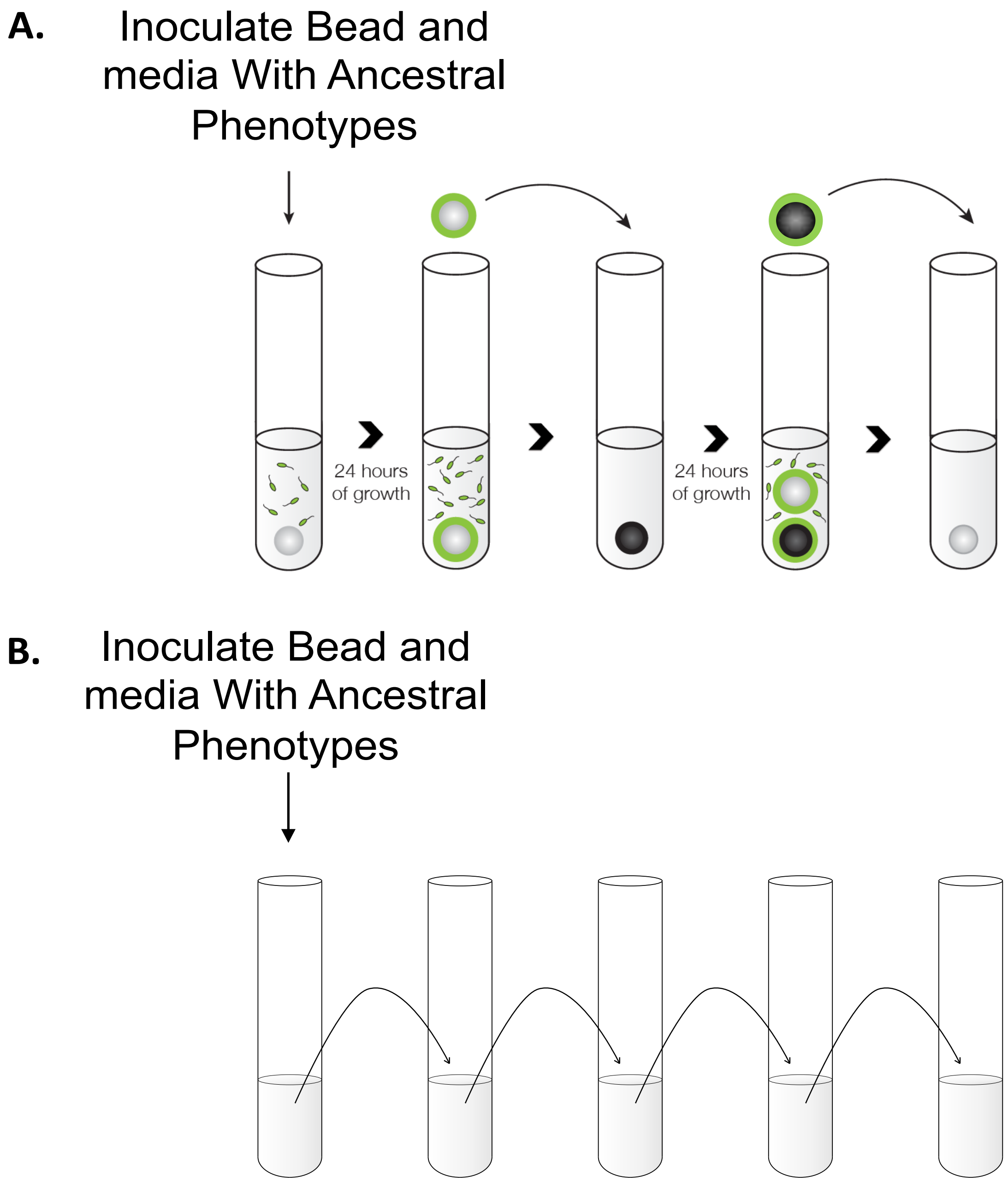


**Figure S1.** Schematic for *Pseudomonas aeruginosa* evolution[10]. (A) Biofilm populations were propagated by transferring on colonized bead to a new tube with an uncolonized bead daily. (B) Planktonic populations were propagated in liquid media with 100-fold dilutions every 24 hours.

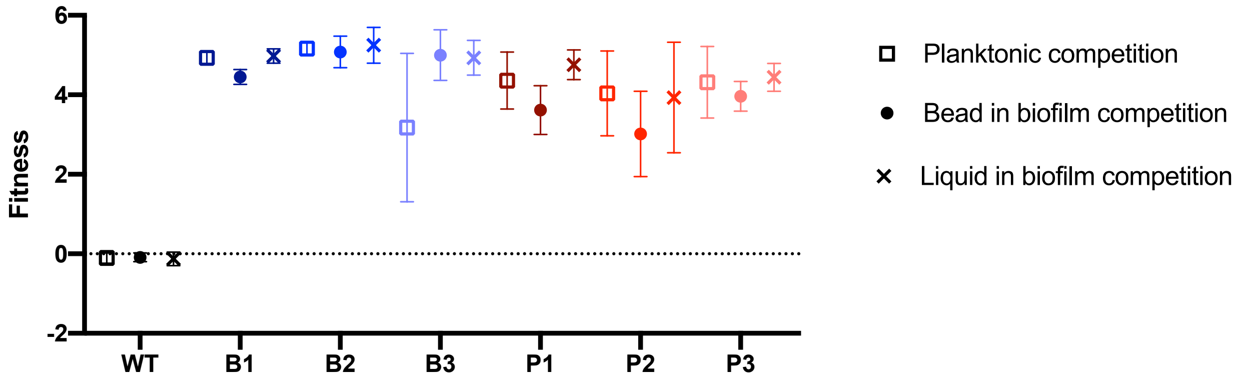


**Figure S2.** 90 day evolved populations have extremely high fitness when competed with the ancestor. Fitness (mean +- 95% confidence interval) of the 90 day populations was determined through pairwise competitions (see methods). Fitness was determined for both planktonic and biofilm competitions. Planktonic competition values are indicated as squares. Biofilm competitions had two measurements: the bead population indicated by filled circles, and the liquid portion of the biofilm assay, indicated by x's. Colors indicate evolved populations with shades of blue indicating biofilm evolved populations, red indicating planktonic evolved populations, and black indicating the ancestral control.

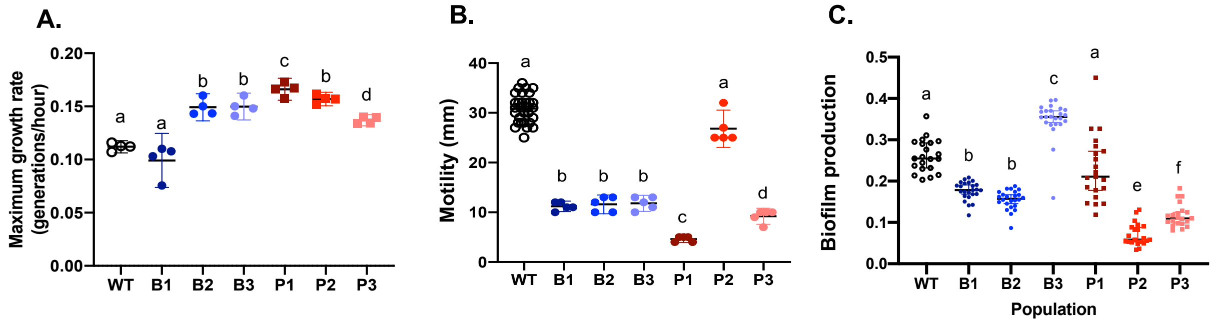


**Figure S3. Traits tied to fitness diversified among replicate populations** . 90 day evolved populations were tested for (A) maximum growth rate; n=4, (B) swimming motility; n = 5, (C) and 4-hr biofilm production; n = 21. All individual populations (B1 = dark blue circles, B2 = blue circles, B3 = light blue circles, P1 = dark red squares, P2 = red squares, P3 = orange squares) were compared against the ancestral, WT, strain (black open circles). All data points are indicated with symbols, with means and 95% CI represented with vertical bars. Letters indicate populations significantly different from one another. There were significant differences observed in all sampled characteristics (ANOVA with Tukey's post hoc testing for A) biofilm production: F=55.17, $p<10^{-4}$, B) motility: F=189.40, $p<10^{-4}$. C) maximum growth rate: F=36.15, $p<10^{-4}$).

**Table S1.** Raw data for fitness calculations. Colony counts are colored by day with counted values under 30 in orange and values of 1 put in for 0's highlighted in yellow.
https://github.com/KatrinaHarris23/PALTEanalysis/blob/master/Table_S1.xlsx

| | All | B1 | B2 | B3 | P1 | P2 | P3 |
|---|---|---|---|---|---|---|---|
| **Mutations:** | 874 | 239 | 180 | 116 | 109 | 94 | 136 |
| **Nucleotide level mutations** | | | | | | | |
| **Indels:** | 31 | 13 | 10 | 2 | 2 | 1 | 3 |
| **Transitions:** | 246 | 107 | 86 | 8 | 6 | 2 | 37 |
| **Transversions:** | 595 | 118 | 84 | 105 | 101 | 91 | 96 |
| **Amino acid level mutations** | | | | | | | |
| **Coding:** | 27 | 11 | 8 | 3 | 2 | 1 | 2 |
| **Intergenic:** | 230 | 53 | 34 | 39 | 37 | 28 | 39 |
| **Pseudogene:** | 10 | 3 | 1 | 4 | 1 | 0 | 1 |
| **Premature stop:** | 9 | 3 | 0 | 2 | 1 | 1 | 2 |
| **Elongating:** | 11 | 4 | 3 | 1 | 2 | 0 | 1 |
| **Synonymous:** | 105 | 33 | 25 | 6 | 16 | 10 | 15 |
| **Non Synonymous:** | 478 | 131 | 108 | 60 | 50 | 53 | 76 |
| **dN/dS:** | 1.75 | 1.53 | 1.66 | 3.85 | 1.20 | 2.04 | 1.95 |

**Table S2. Mutational breakdown for all unique loci mutated in the six evolved populations**. Mutations are broken down at both the nucleotide level and the amino acid level (naming corresponding to breseq output[82]; rows) for all mutations across the study, and looking at each evolved population individually (columns). All unique loci mutated within each population at any time point sequenced are included. dN/dS ratios are standardized to the PA14 genome neutral ratio (see methods).

**Table S3.** All mutational calls determined through whole genome, whole population sequencing of the six populations of PA propagated for 90 days after filtering; see methods for filtering criteria.
https://github.com/KatrinaHarris23/PALTEanalysis/blob/master/Table_S3.csv

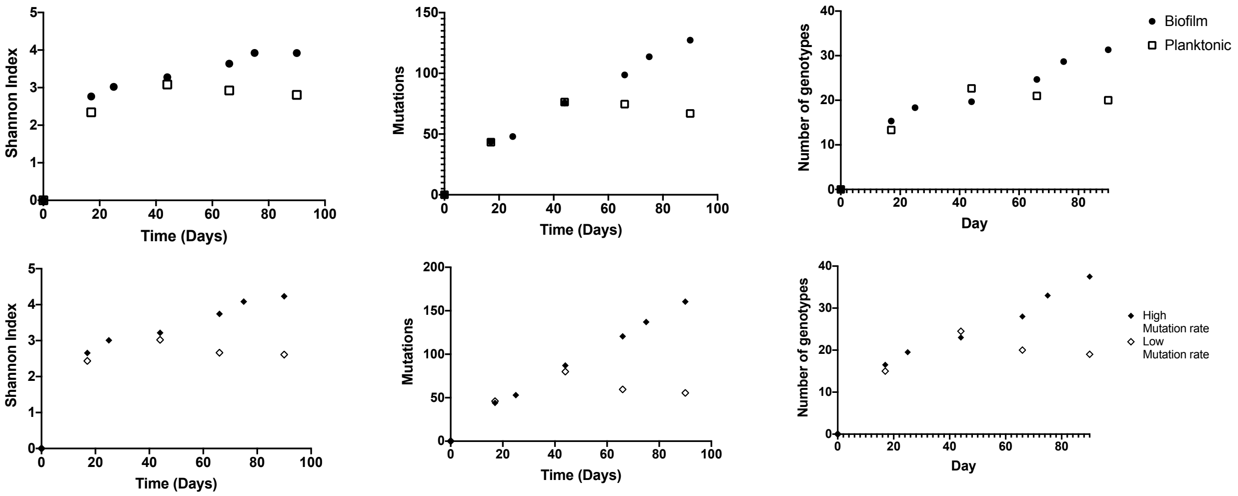


**Figure S4.** Biofilm populations become more diverse over time, in terms of alpha diversity, individual mutations and predicted genotypes plotted through time (top row), but high mutation rate (B1 and B2) compared to low mutation rate (P1 and P2) does not differentiate populations (closed diamonds and open diamonds, respectively; bottom row).

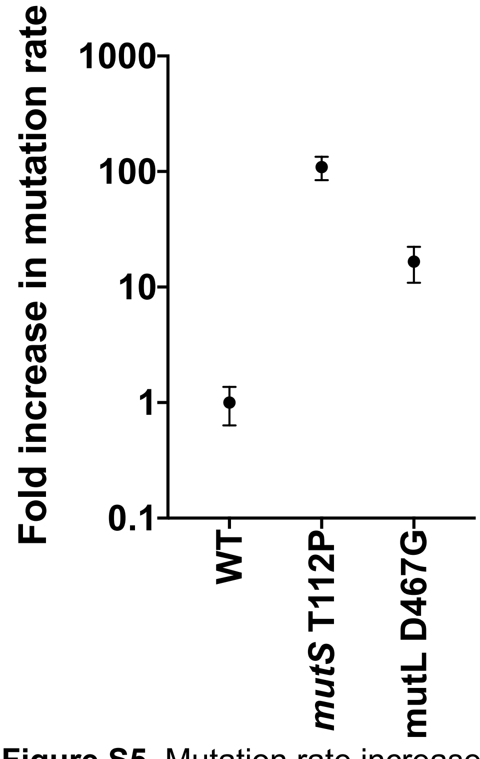


**Figure S5.** Mutation rate increase of *mutS* T112P and *mutL* D467G isogenic mutants. The two mutator alleles that evolved in biofilm environments result in a 109-fold and 16-fold increase in mutation rates over the ancestral strain.

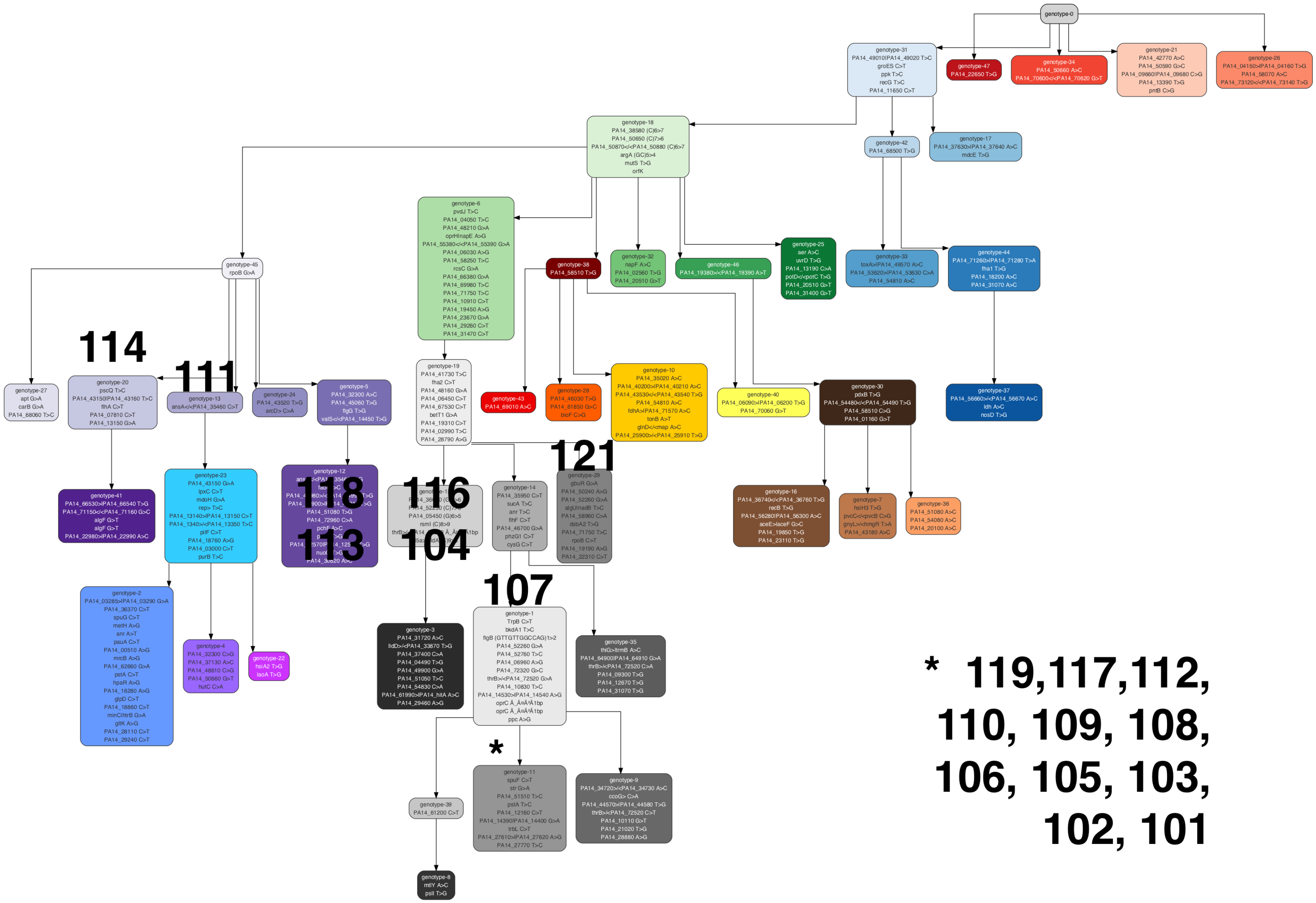


**Figure S6.** Genotype ancestry plot showing the ancestral relatedness of genotypes in population B1. Different colors indicate different genotypes, with shades of the same color indicating the same lineage, with time on the vertical axis. Clones isolated from the B1 population belong to 7 nodes on the B1 population ancestry, indicated by numbers.

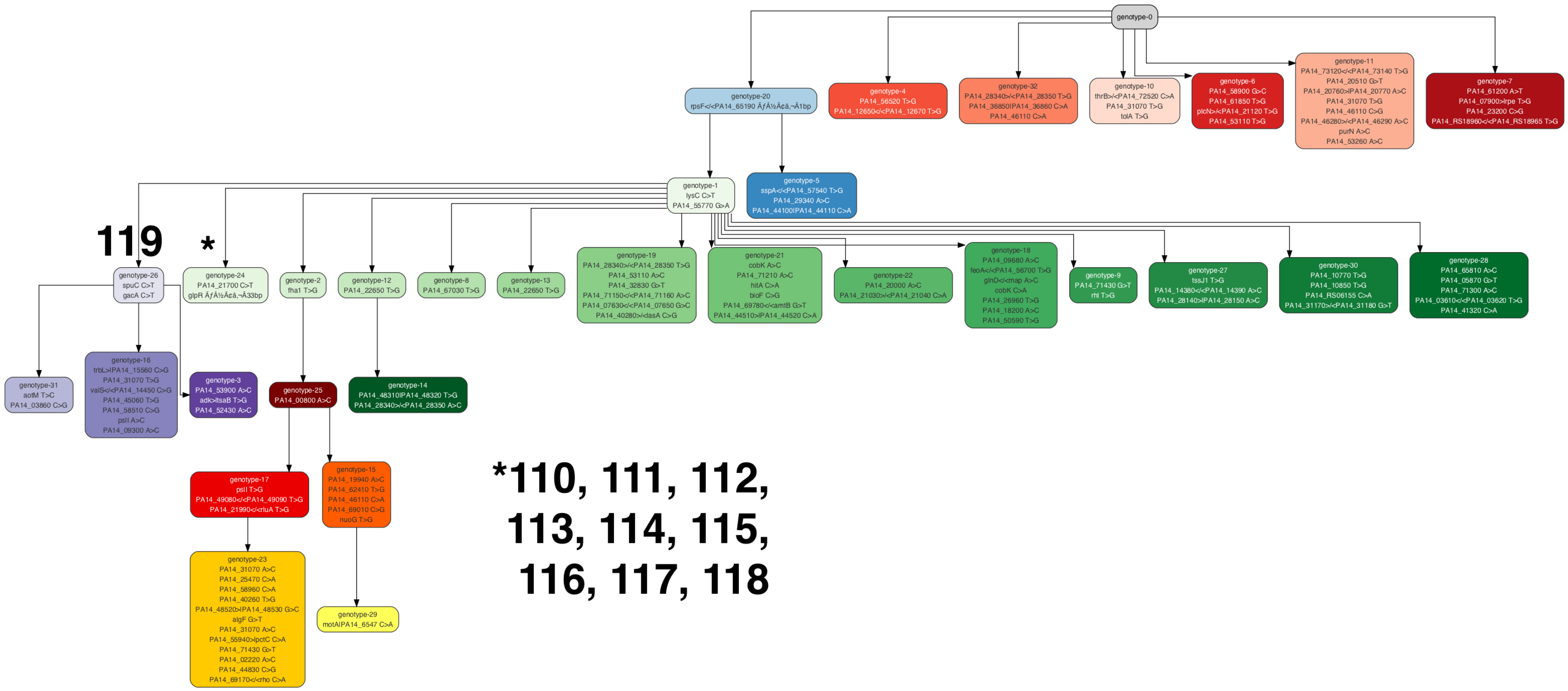


**Figure S7** Clones isolated from the P1 population belong to 2 nodes on the P1 population ancestry.

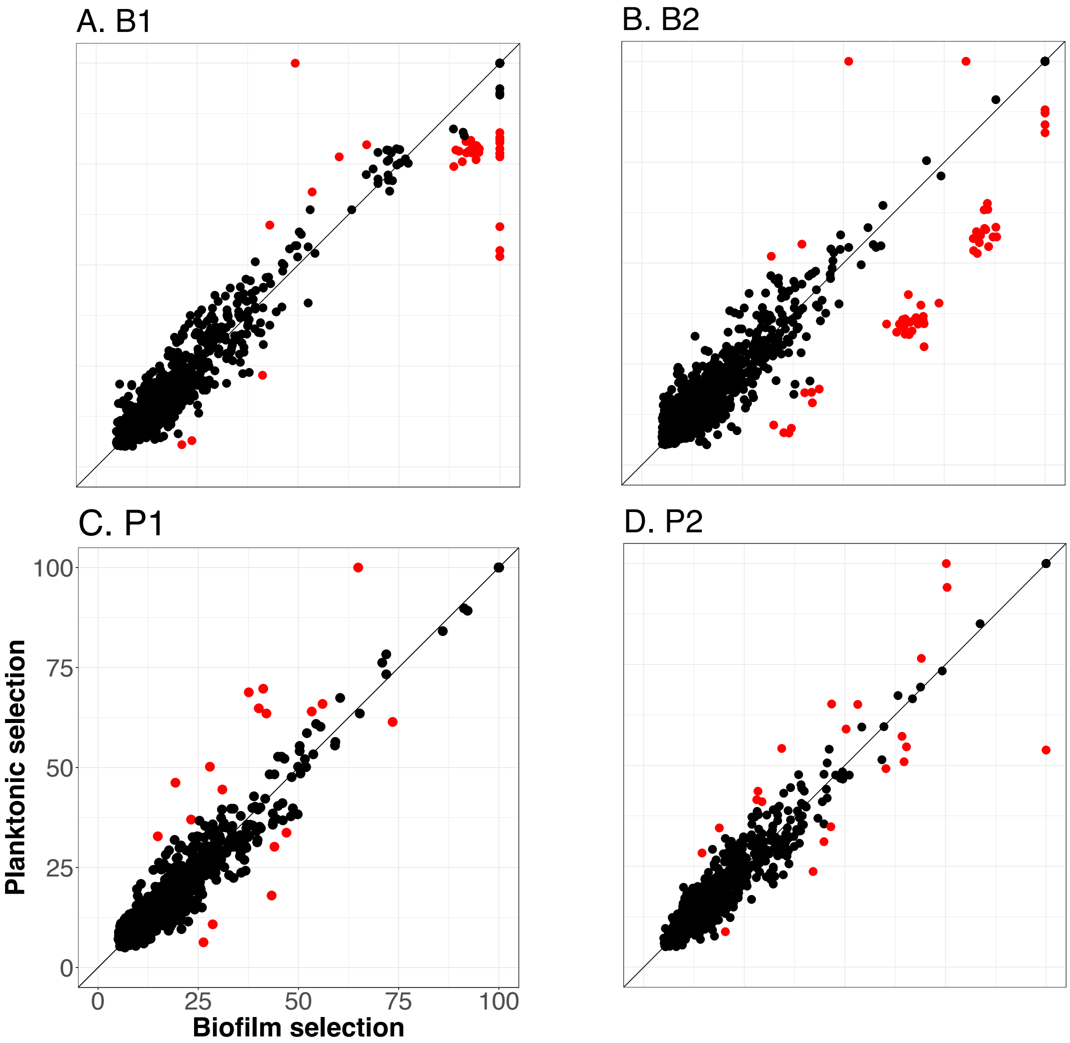


**Figure S8.** Predicted ecological signals are absent from WGPS data after 2 days of selection. Correlation of mutational frequencies when resurrected population DNA is isolated after two days of planktonic selection versus biofilm selection. Analysis was performed on 90-day A. B1 population, B. B2 population, C. P1 population, and D. P2 population. Mutations determined to be significantly enriched through Cook's distance (see methods) are represented in red with nonsignificant mutations in black.

**Table S4.** All mutational frequencies that were determined to be significantly enriched in either biofilm or planktonic conditions during testing for ecological adaptations. Mutations correspond to red points in Fig S8.
https://github.com/KatrinaHarris23/PALTEanalysis/blob/master/Table_S4.xlsx
